# Chemical Crosstalk between the Common Soil Bacteria *Pseudomonas chlororaphis* and *Bacillus subtilis*

**DOI:** 10.1101/2025.07.22.666074

**Authors:** Zohar Tik, Shaked Uzi-Gavrilov, Bat El Hagbi-Lazar, David C. Nwobodo, Orit Sivan, Michael M. Meijler

## Abstract

Chemical conversations between soil inhabitants are of tremendous importance to the health of many ecosystems, and at the same time detailed molecular knowledge underlying these conversations is surprisingly scarce. One of the major bacterial genera inhabiting the rhizosphere is *Pseudomonas*, of which most species are known to produce phenazines, which carry antibiotic properties. *Pseudomonas chlororaphis*, a common rhizosphere dwelling species with plant growth-promoting traits, produces phenazine-1-carboxamide (PCN). This study examines how the production of PCN by *Pseudomonas* affects another common species in soil that it often encounters, namely *Bacillus subtilis*. When both species were cultured at close distance, distinct and visible changes in colony morphologies were observed without changes in growth rates. Interestingly, a clear transformation occurred in the morphology of *B. subtilis* colonies in the presence of supplemented PCN, indicating the role of phenazines in affecting colony morphology. In addition, untargeted metabolomics analyses showed a decrease in the production of plipastatin and surfactin by *B. subtilis* in the presence of *P. chlororaphis*. Our results indicate that PCN induces changes in morphology and signaling of *B. subtilis* without significantly affecting its growth. We hypothesize that *P. chlororaphis* and *B. subtilis* sense one another and act to conserve energy while avoiding competition.

## Introduction

The rhizosphere is the narrow region of soil directly influenced by root secretions and associated soil microorganisms.^1^ As such, the rhizosphere contains higher concentrations of compounds that are secreted by microorganisms and roots than bulk soil,^2^ creating this area a desirable habitat for many microorganisms. Therefore, many intricate microbial interactions occur in the rhizosphere,^3^ creating a fascinating and complicated network of connections between the microorganisms living in the rhizosphere and nearby plants.^2^ Among the rhizosphere microbiome both plant-growth-promoting bacteria (PGPB) and harmful organisms can be found.^4^ PGPBs are beneficial to plants for various reasons, such as promoting nutrient availability^5^ and antibiotic production that protect the plant from pathogens.^6^ Moreover, the diversity of the rhizosphere microbiome allows the plant to adapt to changing environments. Antibiotic production by bacteria is not only taking part in protection against pathogens, but also have a significant effect on the population, and it is a part of its social interaction.^7^ Furthermore, it have a decisive role in the bacteria ability to compete and survive in its surrounding.^7^

*Bacillus* spp., a Gram-positive bacteria, are among the most beneficial PGPB.^8^ A well-studied model organism for this group is the non-pathogenic *Bacillus subtilis*. About 4% of its genome is dedicated to producing specialized metabolites – a relatively high percentage;^9^ this can be indicative of its versatile mechanisms in response to different signals from its neighbors. In *B. subtilis* three different non-ribosomal peptides (NRPs) systems were found, and one hybrid NRP/polyketide.^10^ NRPs distinguish from ribosomal peptides as they contain non-proteinogenic amino acids,^11^ and their structures are quite often macrocyclic.^11,12^ NRPs have diverse biological activities^13^ and are a major chemical family of secondary metabolites.^14^ Several specialized metabolites produced by *B. subtilis* are antibiotics belonging to the family of NRPs. Two of the antibiotics that are produced by those systems are plipastatin and surfactin.^15^ Both plipastatin and surfactin are families of lipopeptides, and the specific molecules within each family differ mainly in the length and saturation of their fatty acid chains.

Another common and well-studied bacterium is the Gram-negative PGPB fluorescent *pseudomonas* spp. *Pseudomonas* is among the most common bacterial genera on earth, and can colonize a wide range of ecological niches, including aquatic areas, plants, animals and soil (i.e. the rhizosphere). *Pseudomonas* are known for their production of phenazines, a family of N-containing heterocyclic compounds, usually pigmented.^16^ Phenazines have important and versatile roles in the soil, as they can function as signaling compounds, antibiotics, electron shuttling and anti-virulence agents.^16–18^ Although this family is highly studied, in most cases their mechanism of action is not fully known.^17^ Not surprisingly, phenazines are known to improve the competitiveness of their producers.^16^ *P. chlororaphis* is a common soil-dwelling species that is considered a prime PGPB.^19–21^ *P. chlororaphis* is known to produce mainly two different phenazines, phenazine-1-carboxamide (PCN) and phenazine-1-carboxylic acid (PCA), and PCN is produced in significantly higher quantities *P. chlororaphis* was found to inhibit the fungal pathogen *Fusarium oxysporum* f. sp. *radicis-lycopersici*, due to its secretion of PCN.^25^ PCN (purified from a different *Pseudomonas* species) was also found to inhibit *Rhizoctonia solani* and *Xanthomonas oryzae* pv. *Oryzae*.^27^

Both *B. subtilis* and *P. chlororaphis* are common in the rhizosphere microbiome, and thus, unsurprisingly, are often co-isolated.^15^ Different interactions were previously reported between different strains from these species, ranging between beneficial, neutral, and harmful.^15^ *B. subtilis* can sense the presence of *P. chlororaphis*, leading to increase in *B. subtilis* sporulation.^28^ Furthermore, *B. subtilis* colonies are protected from infiltration by *P. chlororaphis* due to their extracellular matrix.^28^ Substances in the *B. subtilis* extracellular matrix, such as surfactin, participate in the interaction between them.^27,28^ Although interactions between these two bacteria significantly affect the rhizosphere, the microbiome of the rhizosphere is of course composed of many more bacteria.^30^ It is reasonable to assume that their additional neighbors also affect interactions between *P. chlororaphis* and *B. subtilis*, and vice versa. Moreover, conclusions from culture experiments on the natural environment should not be made easily, as it was established that different growth media and conditions in the lab can change the interactions between those two bacteria.^31^ Nevertheless, further studies on the interactions among those two important PGPBs could extend and deepen our understanding of natural soils.

In this study we examined chemical interactions between *B. subtilis* and *P. chlororaphis*, in order to understand how regulation of these interactions affects their coexistence. We focused on measurable changes in colony morphology, growth and metabolite secretion of both bacteria.

## Results and discussion

### The interaction between *B. subtilis* and *P. chlororaphis* results in colony morphology changes

Wild-type *B. subtilis* strain NCIB 3610 (from here named *B. subtilis*; the microbial strains used in this research are listed in table 1) and wild-type *P. chlororaphis* strain PCL 1391 (from here *P. chlororaphis*) are both prominent members of the rhizosphere; therefore, we set out to examine potentialinteractions between those two species on solid media (agar) in order to best mimic natural conditions. We incubated the bacteria at varying distances (0.4 cm to 1.8 cm) (Figure S1) on LB agar. LB agar was chosen, and not a more selective medium, in order to allow for robust and equal growth of both species. At the shortest distances, 0.4 cm and 0.6 cm, a small inhibition zone in the *B. subtilis* colony was visible, indicating growth inhibition of *B. subtilis* by *P. chlororaphis*. Although this limited inhibition of *B. subtilis* growth occurred in this experiment, the bacteria do often co-exist in nature.^28^ In soil, this inhibition might be prevented due to various reasons such as environmental conditions supporting *B. subtilis* growth, lower concentration of *P. chlororaphis*, and more complicated interactions involving additional bacteria. As our main goal was to study the chemical interactions that actually support their co-existence, we conducted our experiments at 0.8 cm separation between the bacteria, the shortest distance where there was no visible inhibition. Although no inhibition was observed when *B. subtilis* and *P. chlororaphis* were grown on the same plate at 0.8 cm separation, there was a distinct and visible change in the edges of colonies of both bacteria, (Figure 1A) with an extensive change in *B. subtilis* morphology, represented as a change in the wrinkled characteristic morphology,^32^ mainly in the edges of the colonies. This observation is supported by previous studies that established that *P. chlororaphis* has an effect on *B. subtilis* biofilms by activation of the *P. chlororaphis* type IV secretion system (T6SS).^29^ T6SS, via its effectors, causes degradation in the *B. subtilis* peptidoglycan layer,^33^ which forms an important component of the *B. subtilis* cell wall. Previous studies have shown that interactions vary with the specific strain involved,^28^ as well as the bacterial growth medium.^29^ Moreover, there was a visible change in *P. chlororaphis* colony edges. Those morphological changes suggest that the bacteria are sensing one another at this distance. Interestingly, certain interactions between members of those genera were reported by others in diverse conditions.^28,31,34^ Most of those interactions were found to be competitive, while others were found to be beneficial.^28^ Lyng and Kovács suggest that most of the interactions found are more negative than positive due to research methods bias, and not necessarily a reflection of processes in the rhizosphere.^28^ Interestingly, morphological changes were detected all around the colonies, including the farthest zones of the bacteria from one another. It implies that the signals secreted by *P. chlororaphis* are either reaching all of the *B. subtilis* area, either as volatile compound or by diffusion, or the signal is detected by *B. subtilis* near the *P. chlororaphis* colony, and then changes propagate throughout the whole *B. subtilis* colony, as discussed more thoroughly in the next section.

**Table 1.** Microbial strains used in this study.

| Bacterial strain | Description | Source |
| --- | --- | --- |
| *B. subtilis* | Wild type NCIB 3610 | Branda *et al.*<sup>22</sup> |
| *B. subtilis* Δ*pps* | deficient in plipastatin synthesis | Maan *et al.*<sup>23</sup> |
| *B. subtilis* Δ*srfAA* | deficient in surfactin synthesis | López *et al.*<sup>24</sup> |
| *P. chlororaphis* | Wild type PCL 1391 | Chin-A-Woeng *et al.*<sup>25</sup> |
| *P. chlororaphis* Δ*phz* | deficient in phenazine synthesis PCL 119 | Chin-A-Woeng *et al.*<sup>25</sup> |
| *F. oxysporum* | *F. oxysporum* f. sp. *radicis-cucumerinum* | Dror *et al.*<sup>26</sup> |

**Figure 1.**
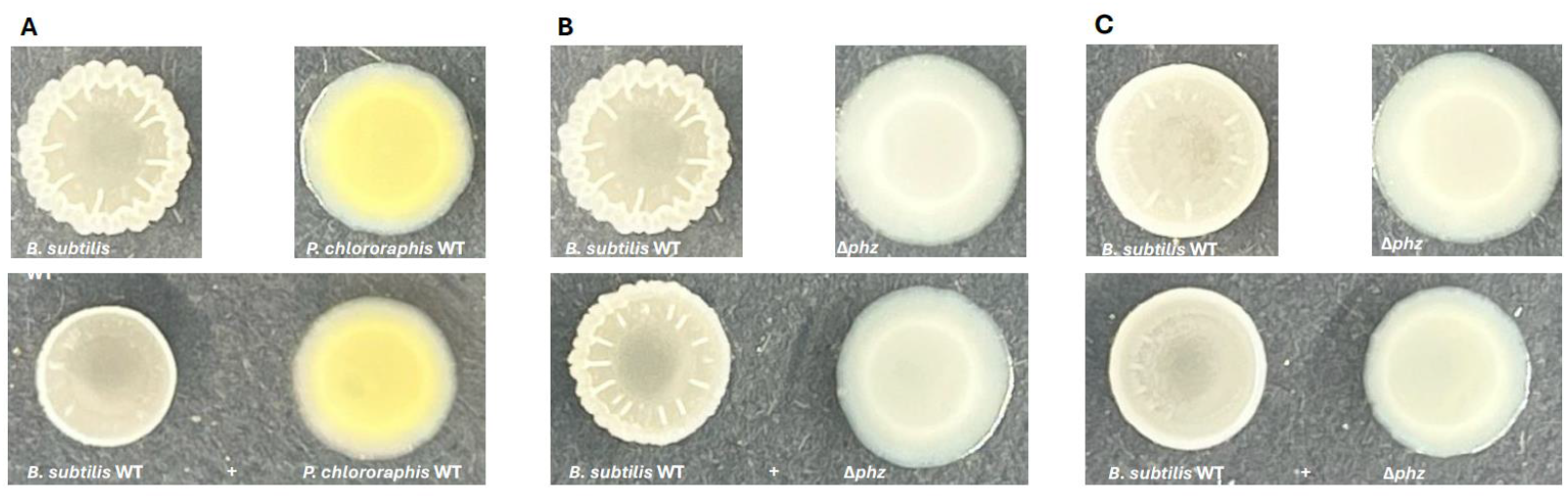
Cultures of *B. subtilis, P. chlororaphis* wild type (WT) and a *P. chlororaphis* non-phenazine producing utant (Δ*phz)*. Top – mono-cultures. Bottom – co-cultures. (A,B) Agar plates (C) Agar + 10 µM PCN plates.

In this study, we examined the potential effects of PCN, the main phenazine produced by *P. chlororaphis*,^35^ on *B. subtilis*, at concentrations that are likely to represent conditions in soil. To further investigate those interactions, we also cultured the *P. chlororaphis* non-phenazine producing mutant strain *P. chlororaphis* 1119 (from here *P. chlororaphis* Δ*phz*). We cultured *B. subtilis* and *P. chlororaphis* Δ*phz* as mono- and co-cultures (Figure 1B), similarly as the wild-type co-cultures. Phenazines are known as colorful compounds, and PCN has a yellow color, a difference that is evident in the *P. chlororaphis* Δ*phz* cultures (Figure 1A,B). However, when *B. subtilis* grew next to *P. chlororaphis* Δ*phz*, there was still a change in the bacterial morphology, although the change was smaller compared to *B. subtilis*-*P. chlororaphis* co-cultures (Figure 1A,B). Not surprisingly, this suggests, that while PCN clearly affects *B. subtilis* morphology, it is not the only compound produced by *P. chlororaphis* that affects *B. subtilis*. This correlates with previous studies showing that other compounds are interacting with *Bacillus* and *Pseudomonas* species, such as 2,4-diacetylphloroglucinol,^36^ and lipopeptides.^37^

### PCN has a significant role in the interaction between *B. subtilis* and *P. chlororaphis*

The clear observation of the differential reaction of *B. subtilis* towards *P. chlororaphis* and *P. chlororaphis* Δ*phz* indicates the role of PCN in the interaction among these two species. Other groups have reported previously that PCN can strongly inhibit the growth of many microorganisms, including fungi, and engineered strains of *P. chlororaphis* that produce high concentrations of PCN are used as biocontrol agents. However, these effects are only observed at higher PCN concentrations (>50 mM, often reaching even 1000 mM), far greater than the low-micromolar PCN concentrations produced by wild-type *P. chlororaphis* strains. Interestingly, a recent study by Zhou et al., that was published in the course of our manuscript submission, identified the target of PCN (from the wheat-associated *P. chlororaphis* strain ZJU60, a high-level PCN producer) in *B. subtilis* strain 3610, as topoisomerase IV.^38^ Previously, the same group reported that PCN from ZJU60 reduces virulence of the fungal plant pathogen *Fusarium graminearum* by altering histone acetylation. However, in both cases only the effects of high concentrations of PCN were studied.^39^

To gain a better understanding of the role that PCN plays in guiding *B. subtilis* and *P. chlororaphis* interactions that are less antagonistic and that closer resemble natural conditions, we examined whether addition of 10 µM PCN to agar would restore the activity of *P. chlororaphis* Δ*phz* to the level of its wild-type counterpart. The PCN concentration was determined to be 10 µM based on PCN concentrations produced by *P. chlororaphis* (Figure S2). Co-cultures with the Δ*phz* mutant on media with PCN displayed similar effects on *B. subtilis* morphology as a co-culture with wild-type *P. chlororaphis*, indicating that changes in *B. subtilis* morphology are dependent on the presence of PCN. (Figure 1A,C, Figure S3). For further discussion of the spatial effect of PCN on *B. subtilis* morphology, see the supporting information (Figure S4).

### The interaction between *B. subtilis* and *P. chlororaphis* results in *B. subtilis* metabolic changes

In order to further our understanding of how the two soil bacteria affect one another, and in particular how PCN affects *B. subtilis* behavior, we set out to examine differences in production of *B. subtilis* metabolites in the presence of *P. chlororaphis* and PCN alone on solid media, and compared with effects caused by the presence of the PCN mutant. To start our analysis, first a principal component analysis by MetaboAnalyst^40^ based on MS^1^ data showed that each treatment clustered by itself (Figure 2A). This result indicates that clear changes in overall metabolite production between the different experimental treatments could be observed. *B. subtilis* grown together with *P. chlororaphis* Δ*phz* was more similar to *B. subtilis* grown on mono-culture than *B. subtilis* co-culturing with *P. chlororaphis*. Moreover, *B. subtilis* cultured together with *P. chlororaphis* Δ*phz* was more similar to *B. subtilis* rather than when cultured together with PCN, indicating a more prominent effect on *B. subtilis* metabolites when the phenazine was present. This observation supports our hypothesis that phenazines alter *B. subtilis* metabolite production. In addition, it should be considered that the metabolic profile of the *P. chlororaphis* Δ*phz* culture with PCN is similar to *P. chlororaphis* Δ*phz* rather *than P. chlororaphis*. The *P. chlororaphis* Δ*phz* mutant used in this study has a transposon insertion at the PhzB homolog gene,^25^ resulting in obstruction of the bacterial phenazine production pathway. That means that *P. chlororaphis* and *P. chlororaphis* Δ*phz* are different in more metabolites than only PCN, which can explain the differences between the mono-cultures treatments shown in figure 2A. After establishing differences in the metabolic profiles, we continued investigating which *B. subtilis* metabolites are the most affected by PCN.

**figure 2.**
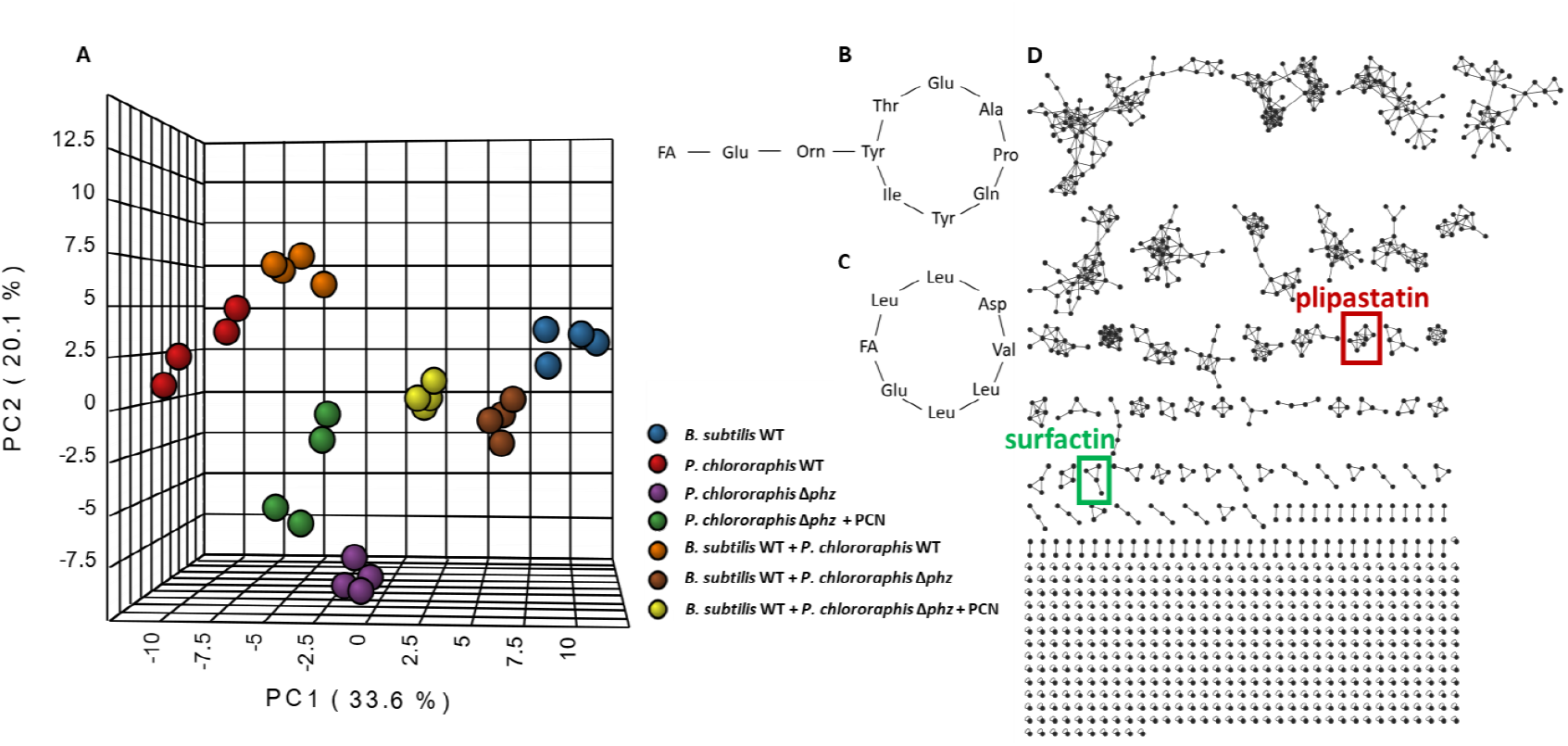
Effects on metabolite secretion in the bacterial co-cultures. (A) Principal component analysis of MS^1^ data from single and co-culture extracts, created with use of MetaboAnalyst. The data is based on four replicates of cultures. Structure of (B) plipastatin and (C) surfactin. (D) Based on MS^2^ data obtained from extracts of the various co-cultures and data treatment using MZmine, a network could be generated using the feature-based molecular network workflow (Nothias et al. 2020) on the open source GNPS platform (Wang et al. 2016). Each node represents a metabolite in the sample, and any edge connecting two nodes suggests that they have a similar structure. Plipastatin and surfactin clusters are indicated in red and green, respectively. FA = fatty acid. For the specific fatty acids identified in this study, see Figures S6, S7.

Several metabolites were found to be produced in altered amounts among the different treatments. We used several metabolomics platforms, MZmine3,^41^ GNPS^42^ and Sirius,^43^ to annotate those affected metabolites; the molecular network produced obtained via the GNPS platform is shown in figure 2D. We aimed to explore compounds that were altered as a result of the different treatments. Although metabolite annotation has progressed much in recent years, unequivocal annotation with a high degree of confidence is still a major bottleneck in untargeted metabolomics research.^44^ Thus, as was expected, we were not yet able to identify all the compounds of interest. For example, we found a compound with an m/z value of 227.1754, which production was significantly inhibited in the presence of *P. chlororaphis*, and also upon the addition of PCN, but its production was not inhibited when *B. subtilis* and *P. chlororaphis* Δ*phz* were co-cultured (Figure S5). The structure with Sirius’s highest annotation score for this mass was the cyclo-di-peptide cyclo-Leu-Leu. Although we did not validate the structure of this compound, we will continue to examine this possibility, as cyclo-di peptides are known to participate in quorum sensing processes and have antimicrobial and antifungal properties.^45^ Furthermore, cyclo-di-peptides were isolated from *Bacillus* spp.^46,47^ Thus, it is not unlikely that a compound from this family can serve to mediate interactions between *B. subtilis* and *P. chlororaphis*. Thus, it appears that the secretion of PCN by *P. chlororaphis* results in inhibition of this compound with unknown structure. However, some of the metabolites whose concentrations were changed due to PCN addition could be annotated by GNPS, and, interestingly, several of these are known *Bacillus*-produced antimicrobial substances, such as plipastatin and surfactin (Figure 2B, 2C, 3A, 3B).

The presence of *P. chlororaphis* results in strongly decreased plipastatin quantities secreted by *B. subtilis*, while the presence of *P. chlororaphis* Δ*phz* does not significantly affect the production of plipastatin by *B. subtilis* (Figure 3A). Furthermore, when *B. subtilis* was grown on PCN-containing media, both alone and together with *P. chlororaphis* Δ*phz*, a similar decrease in plipastatin production was observed (Figure 3A), implying PCN involvement in a mechanism lowering this antimicrobial substance production in *B. subtilis*. The same effect was detected for other plipastatin derivatives (supporting information). Similarly, the production of surfactin, another antimicrobial compound produced by *B. subtilis*, is strongly inhibited by the presence of PCN, although in the co-culture experiments the differences were smaller (Figure 3B), indicating the involvement of additional signals of *P. chlororaphis* that may antagonize the effects of PCN on *B. subtilis*. It was previously suggested that surfactin is involved in the interactions between those two species,^29^ as we observed as well in this study, and it appears that different mechanisms tightly control the production of surfactin. *B. subtilis* was shown to decrease the production of both plipastatin and surfactin when interacting with the fungus *Setophoma terrestris*.^48^ The interaction between *B. subtilis* and *S. terrestris* resulted in growth inhibition of the fungus only when exposed to *B. subtilis* cell-free supernatant after exposure of *B. subtilis* to *S. terrestris*.^49^

**Figure 3.**
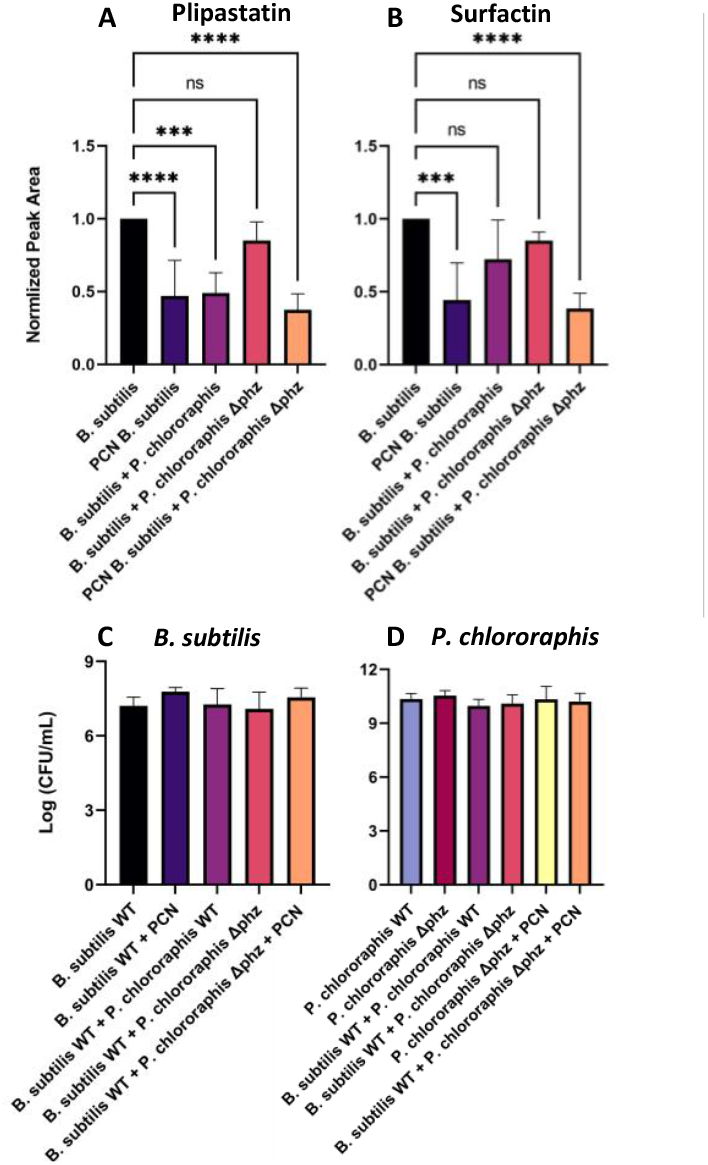
Production of (**A**) plipastatin, and (**B**) surfactin by *B. subtilis* in various treatments. The data represent the average of normalized peak area of three biological replicates by *B. subtilis* secretion in mono-culture, consist of four technical replicates, based on relevant compound production in mono-culture of *B. subtilis*. Colony forming unit of (**C**) *B. subtilis*, and (**D**) *P. chlororaphis* WT and *P. chlororaphis* Δ*phz*. Statistical analysis was performed using one-way ANOVA, with p<0.0001, p<0.001, are marked by ****, ***, respectively.

To verify that the decrease in *B. subtilis* antibiotic production was not caused by inhibition of the bacterial growth, we analyzed growth (by counting CFUs) in the various treatments (Figure 3C, D). Both *B. subtilis* and *P. chlororaphis* (WT and Δ*phz*) did not show a significant change in their CFU count while cultured together, or in treatments where PCN was added. Moreover, we cultured *B. subtilis*, its non-plipastatin-producing mutant, *B. subtilis* Δ*pps*, and its non-surfactin-producing mutant, *B. subtilis* Δ*srfAA*, in the presence of different PCN concentrations (Figure 4). A similar trend was observed among the different bacterial strains. At the highest PCN concentrations examined, 32.3-250 µM, all three types of bacteria were inhibited (Figure 4). Although these results are exciting, especially given the general importance of such phenazines, further studies are needed to determine their concentrations, and thus relevance to microbial growth inhibition, in natural environments.^50^ PCA, an additional phenazine abundant in *Pseudomonas* species, is more comprehensively studied than PCN. Mavrodi et al.^50^ extracted PCA with the highest concentration of 1600 ng g^-1^ of roots (approximately 7.1 µM) in a different study on various phenazine-producing (Phz^+^) *Pseudomonas* spp However, further studies are warranted to determine accurate concentrations of PCN and other phenazines in natural environments.^50^ At 15.6 µM of PCN, a slight growth inhibition was shown only in *B. subtilis* Δ*srfAA* towards the end of the experiment (Fig. 4C). At lower PCN concentrations, there was no difference between the treatments and the control that did not contain PCN. This indicates that at medium-low concentrations of PCN, which are naturally relevant, plipastatin and surfactin have no major part in *B. subtilis* protection mechanism against PCN.

**Figure 4.**
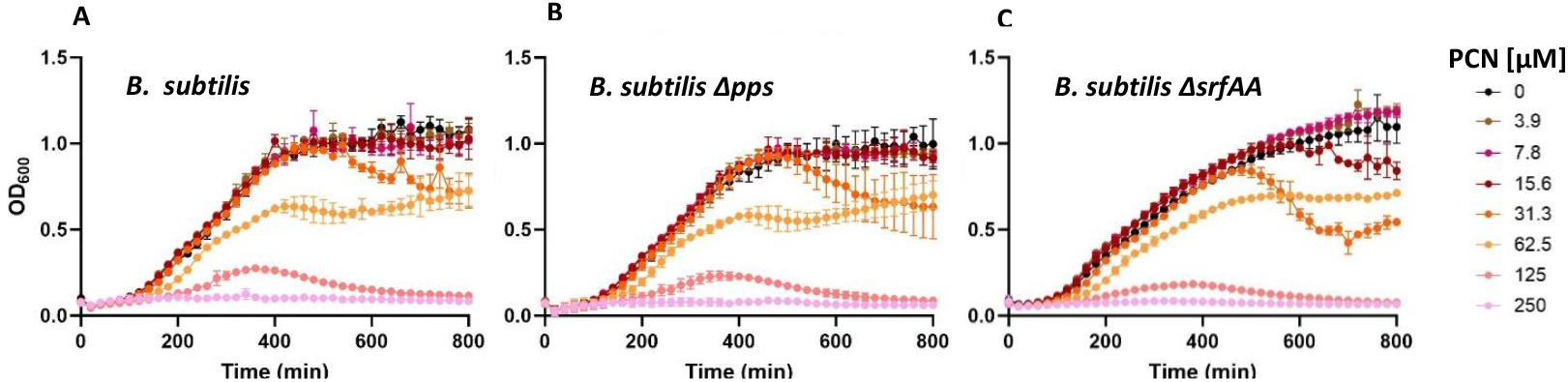
PCN affects *B. subtilis* growth: (A) WT (B) Δ*pps* (C) Δ*srfAA*. Error bars represent the standard deviation of triplicates and are not shown when smaller than the symbol.

### Plipastatin and surfactin affect *P. chlororaphis* phenotypes

Given that the presence of *P. chlororaphis*, through secretion of PCN, results in a decrease of plipastatin and surfactin production in *B. subtilis* (Figure 3), we propose two hypotheses to explain this phenomenon: (i) competitive interactions: as *B. subtilis* antibiotics such as surfactin and plipastatin inhibit the growth of *P. chlororaphis*, the latter may have evolved to inhibit the production of those compounds. However, *B. subtilis* could potentially have evolved to overcome this inhibition through upregulation of anti-biotics production upon sensing PCN, which we did not observe; (ii) this leads us to propose that we may observe here cooperative interactions: the presence of *P. chlororaphis* antibiotics (that do not inhibit *B. subtilis* significantly) in the environment may trigger a sensing mechanism in *B. subtilis*, that leads to downregulation of antibiotic production pathways, thereby effectively conserving energy by avoiding redundant antibiotic synthesis, as it can coexist with *P. chlororaphis*.

To investigate these hypotheses further, we examined differences in metabolites produced by *P. chlororaphis* while growing adjacent to *B. subtilis* or *B. subtilis* Δ*pps*. We observed no significant differences in the production of *P. chlororaphis* metabolites. To further examine the mechanism underlying decreased *B. subtilis* antibiotics production, we isolated a derivative of plipastatin, plipastatin A C_19_ (Figure S6), from *B. subtilis* cultures, along with a mixture of surfactins (Figure S7). We incubated *P. chlororaphis* with increasing concentrations of these isolated metabolites. We did not observe growth inhibition in the presence of these metabolites, in fact we did observe a slight increase in *P. chlororaphis* growth, with surfactin displaying a stronger effect than plipastatin (Figures 5A, B, S8). As a result, we assume that additional signals (and not only PCN) are involved in the interactions between the bacteria.

**Figure 5.**
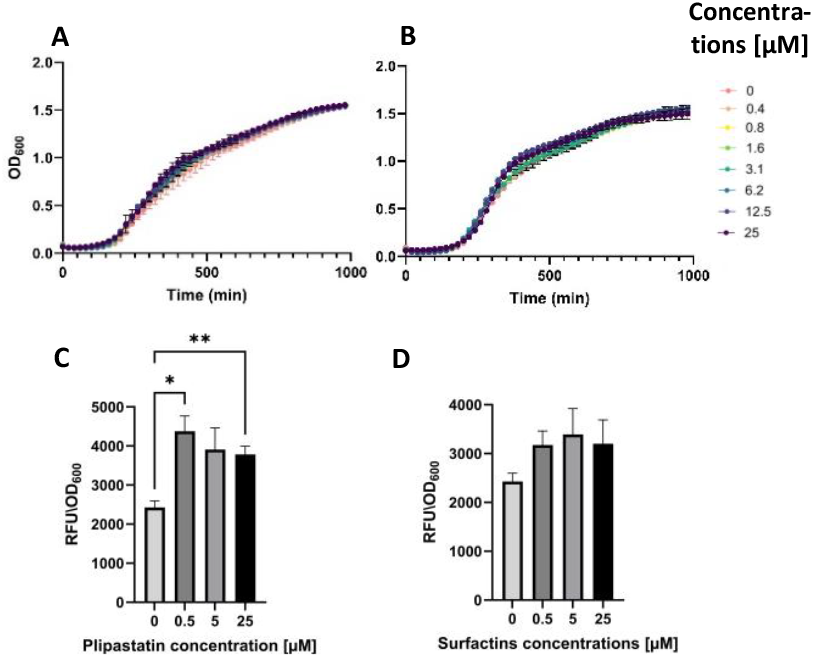
Effects of plipastatin and surfactin on *P. chlororaphis* growth (A, B) and ROS production (C, D). As test compounds we examined plipastatin A C_19_ (A, C) and a mixture of surfactins (B, D). Error bars represent the standard deviation of triplicates. Error bars smaller than the symbol are not shown. Statistical analysis was performed using Brown-For-sythe and Welch ANOVA, with P<0.01 and p<0.05 marked by **, *, respectively.

To examine additional interactions between the bacteria and the effects of plipastatin and surfactin on *P. chlororaphis*, we measured the production of reactive oxygen species (ROS) by the bacteria in the presence of the different *Bacillus* antibiotics. ROS are a group of highly reactive molecules derived from molecular oxygen. Bacteria can produce ROS and inhibit other microorganisms in their surroundings, and *Pseudomonas* spp. are especially adept at their production.^51,52^ To quantify the ROS production by *P. chlororaphis*, we used 2′,7′-dichlorodihydrofluorescein diacetate (DCFDA). Upon exposure to ROS, DCFDA undergoes oxidation to 2’,7’-dichlorofluorescein (DCF), which results in an increase of fluorescence. The addition of both plipastatin and surfactins to the bacteria resulted in an increase in ROS production (Figure 5C, D), in line with previous studies demonstrating that *B. subtilis* employs plipastatin as a defense strategy against the plant pathogens *Magnaporthe grisea*^53^ and *Rhizopus stolonifer*.^54^ This mechanism involves enhancing ROS production, which contributes to the antifungal activity of *B. subtilis*. However, our results show that these increases do not appear to significantly limit the growth of *B. subtilis* (Figure 3C). We propose, however, that ROS production by *P. chlororaphis* could benefit both species by creating an environment that is less attractive for other species to establish colonies, and *B. subtilis* has effective tools to cope with ROS.^55^

Our initial observations suggest that ROS production by *P. chlororaphis* may provide a communal benefit by creating an environment that is less conducive to colonization by other microbial species, potentially favoring the persistence of both *P. chlororaphis* and *B. subtilis*. To further examine this hypothesis, we evaluated the effect of cell-free supernatants (CFS) from both monocultures and cocultures on the growth of the soil-borne fungal plant pathogen *Fusarium oxysporum*. Interestingly, CFS from monocultures of each bacterium exhibited higher antifungal activity, whereas coculture CFS showed comparatively reduced inhibition (Figures S9-S12). This observation aligns with the hypothesis that interspecies sensing between *P. chlororaphis* and *B. subtilis* may trigger a strategy of energy conservation, wherein both bacteria downregulate costly secondary metabolite production in the presence of a partner species. Although in our laboratory conditions *B. subtilis* growth was somewhat inhibited in cocultures (Figure S12), *F. oxysporum* growth was inhibted strongly when incubated with both *P. chlororaphis* monocultures and the cocultures (Figures S9-S12). Concurrently, ROS production by *P. chlororaphis* may contribute to establishing a mildly hostile environment, reducing the likelihood of colonization by third-party competitors. Together, these mechanisms suggest that coculture interactions balance metabolic expenditure with ecological defense, optimizing survival while limiting unnecessary energetic costs associated with antagonistic metabolite production. We examined thus far only one of many potential fungal soil coinhabitants that these bacteria often encounter, and these effects may well differ when other fungi will be examined, which is part of our ongoing studies.

Taken together, our observations support the hypothesis that co-evolution of *B. subtilis* and *P. chlororaphis* may have led to mutualistic rather than competitive interactions, through down-regulation of antibiotic agents such as plipastatin and surfactins, as depicted in Figure 6.

**Figure 6.**
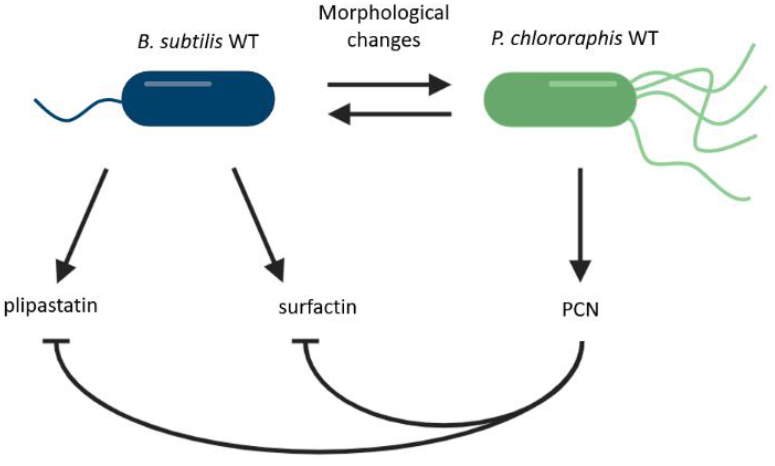
Figure 6. Proposed chemical interactions between *B. subtilis* (NCIB 3610) and *P. chlororaphis* (PCL 1391) and their effects.

## Conclusions

*B. subtilis* and *P. chlororaphis* evolved together and interact with one another. Our study reveals that while the interaction seems to alter production of *B. subtilis* metabolites, it is not the case for *P. chlororaphis*. However, morphological changes were observed in both species. This interaction resulted in lower antibiotic production by *B. subtilis*. Our results support the hypothesis that lowering antibiotic production is part of a cooperation mechanism, not a competititive one. The production of plipastatin and surfactins is costly for *B. subtilis*, and as it appears to be able to cope with physiological concentrations of PCN (up to 15 µM), it may be advantageous for *B. subtilis* to lower its antibiotic production and allow for the effects of *P. chlororaphis* virulence factors to provide a ‘protective zone’. Nevertheless, environmental interactions are obviously much more complex than interactions under controlled laboratory conditions, examining only a few species. Further research is needed to examine those interactions under conditions resembling those of the natural environment, such as addition of specific soil minerals and additional fungal species.

## Supporting information

Supporting information including experimental procedures and figures S1-S12

## ASSOCIATED CONTENT

Experimental procedures and supporting figures 1–12, including additional experiment photos, metabolites distributions and mass spectrometry chromatograms.

The MS data are accessible at the UCSD Center for Computational Mass Spectrometry’s server (access with password) which will become public upon publication.

## AUTHOR INFORMATION

### Authors

Shaked Uzi-Gavrilov^‡^ - *Department of Chemistry, Ben-Gurion University of the Negev, 8410501, Be’er Sheva, Israel*.

Bat El Hagbi-Lazar - *Department of Chemistry, Ben-Gurion University of the Negev, 8410501, Be’er Sheva, Israel*.

David C. Nwobodo - *Department of Chemistry, Ben-Gurion University of the Negev, 8410501, Be’er Sheva, Israel*

Orit Sivan - *Department of Earth and Environmental Sciences, Ben-Gurion University of the Negev, 8410501, Be’er Sheva, Israel*.

### Present Addresses

†Zohar Tik – Department of Plant and Environmental Sciences, Weizmann Institute of Science, 7610001, Rehovot, Israel.

^‡^Shaked Uzi-Gavrilov – Department of Plant and Microbial Biology, North Carolina State University, 27695, North Carolina, United States.

## Author Contributions

All authors have given approval to the final version of the manuscript.

## ACKNOWLEDGMENT

We would like to thank Dianne K. Newman and her group members for providing *P. chlororaphis* strains and for helpful discussions and assistance in this research. Ilana Kolodkin-Gal kindly provided us with *B. subtilis* strains used in this study. Omer Frenkel kindly provided us with *Fusarium oxysporum*.

## REFERENCES

(1) Philippot, L.; Raaijmakers, J. M.; Lemanceau, P.; Van Der Putten, W. H. Going Back to the Roots: The Microbial Ecology of the Rhizosphere. Nat. Rev. Microbiol. 2013 1111 2013, 11 (11), 789–799. 10.1038/nrmicro3109.

(2) Turner, T. R.; James, E. K.; Poole, P. S. The Plant Microbiome. Genome Biol. 2013, 14 (6), 1–10. 10.1186/gb-2013-14-6-209.

(3) Minz, D.; Ofek, M. Rhizosphere Microorganisms. Benef. Microorg. Multicell. Life Forms 2012, 105–121. 10.1007/978-3-642-21680-0_7.

(4) Yu, K.; Pieterse, C. M. J.; Bakker, P. A. H. M.; Berendsen, R. L. Beneficial Microbes Going Underground of Root Immunity. Plant. Cell Environ. 2019, 42 (10), 2860–2870. 10.1111/PCE.13632.

(5) Etesami, H.; Adl, S. M. Plant Growth-Promoting Rhizobacteria (PGPR) and Their Action Mechanisms in Availability of Nutrients to Plants. 2020, 147–203. 10.1007/978-981-15-2576-6_9.

(6) Saeed, Q.; Xiukang, W.; Haider, F. U.; Kučerik, J.; Mumtaz, M. Z.; Holatko, J.; Naseem, M.; Kintl, A.; Ejaz, M.; Naveed, M.; Brtnicky, M.; Mustafa, A. Rhizosphere Bacteria in Plant Growth Promotion, Biocontrol, and Bioremediation of Contaminated Sites: A Comprehensive Review of Effects and Mechanisms. Int. J. Mol. Sci. 2021, Vol. 22, Page 10529 2021, 22 (19), 10529. 10.3390/IJMS221910529.

(7) Cordero, O. X.; Wildschutte, H.; Kirkup, B.; Proehl, S.; Ngo, L.; Hussain, F.; Le Roux, F.; Mincer, T.; Polz, M. F. Ecological Populations of Bacteria Act as Socially Cohesive Units of Antibiotic Production and Resistance. Science (80-.). 2012, 337 (6099), 1228–1231. 10.1126/science.1219385.

(8) Sansinenea, E. Bacillus Spp.: As Plant Growth-Promoting Bacteria. Second. Metab. Plant Growth Promot. Rhizomicroorganisms Discov. Appl. 2019, 225–237. 10.1007/978-981-13-5862-3_11.

(9) Earl, A. M.; Losick, R.; Kolter, R. Ecology and Genomics of Bacillus Subtilis. Trends Microbiol. 2008, 16 (6), 269–275. 10.1016/J.TIM.2008.03.004.

(10) Hou, Q.; Kolodkin-Gal, I. Harvesting the Complex Pathways of Antibiotic Production and Resistance of Soil Bacilli for Optimizing Plant Microbiome. FEMS Microbiol. Ecol. 2020, 96 (9), 142. 10.1093/FEMSEC/FIAA142.

(11) Schwarzer, D.; Finking, R.; Marahiel, M. A. Nonribosomal Peptides: From Genes to Products. Nat. Prod. Rep. 2003, 20 (3), 275–287. 10.1039/B111145K.

(12) Finking, R.; Marahiel, M. A. Biosynthesis of Nonribosomal Peptides. Annu. Rev. Microbiol. 2004, 58 (Volume 58, 2004), 453–488. 10.1146/annurev.micro.58.030603.123615.

(13) Comert Onder, F.; Ay, M. Recent Developments in Natural Bioactive Peptides: Anticancer Potential and Structure–Activity Relationships. Stud. Nat. Prod. Chem. 2022, 75, 329–365. 10.1016/B978-0-323-91250-1.00009.

(14) Duban, M.; Cociancich, S.; Leclère, V. Nonribosomal Peptide Synthesis Definitely Working Out of the Rules. Microorg. 2022, Vol. 10, Page 577 2022, 10 (3), 577. 10.3390/MICROORGANISMS10030577.

(15) Tran, C.; Cock, I. E.; Chen, X.; Feng, Y. Antimicrobial Bacillus: Metabolites and Their Mode of Action. Antibiotics 2022, 11 (1), 88. 10.3390/ANTIBIOTICS11010088.

(16) Mavrodi, D. V.; Blankenfeldt, W.; Thomashow, L. S. Phenazine Compounds in Fluorescent Pseudomonas Spp. Biosynthesis and Regulation*. Annu. Rev. Phytopathol. 2006, 44, 417–445. 10.1146/ANNUREV.PHYTO.44.013106.145710.

(17) Biessy, A.; Filion, M. Phenazines in Plant-Beneficial Pseudomonas Spp.: Biosynthesis, Regulation, Function and Genomics. Environ. Microbiol. 2018, 20 (11), 3905–3917. 10.1111/1462-2920.14395.

(18) Price-Whelan, A.; Dietrich, L. E. P.; Newman, D. K. Rethinking “secondary” Metabolism: Physiological Roles for Phenazine Antibiotics. Nat. Chem. Biol. 2006 22 2006, 2 (2), 71–78. 10.1038/nchembio764.

(19) Shen, X.; Chen, M.; Hu, H.; Wang, W.; Peng, H.; Xu, P.; Zhang, X. Genome Sequence of Pseudomonas Chlororaphis GP72, a Root-Colonizing Biocontrol Strain. J. Bacteriol. 2012, 194 (5), 1269–1270. 10.1128/JB.06713-11.

(20) Brilli, F.; Pollastri, S.; Raio, A.; Baraldi, R.; Neri, L.; Bartolini, P.; Podda, A.; Loreto, F.; Maserti, B. E.; Balestrini, R. Root Colonization by Pseudomonas Chlororaphis Primes Tomato (Lycopersicum Esculentum) Plants for Enhanced Tolerance to Water Stress. J. Plant Physiol. 2019, 232, 82–93. 10.1016/J.JPLPH.2018.10.029.

(21) Miguelez-Sierra, Y.; Acebo-Guerrero, Y.; El Jaziri, M.; Bertin, P.; Hernández-Rodríguez, A. Pseudomonas Chlororaphis CP07 Strain Reduces Disease Severity Caused by Phytophthora Palmivora in Genotypes of Theobroma Cacao. Eur. J. Plant Pathol. 2019, 155 (4), 1133–1143. 10.1007/s10658-019-01842-8.

(22) Branda, S. S.; González-Pastor, J. E.; Ben-Yehuda, S.; Losick, R.; Kolter, R. Fruiting Body Formation by Bacillus Subtilis. Proc. Natl. Acad. Sci. U. S. A. 2001, 98 (20), 11621–11626. 10.1073/pnas.191384198.

(23) Maan, H.; Itkin, M.; Malitsky, S.; Friedman, J.; Kolodkin-Gal, I. Resolving the Conflict between Antibiotic Production and Rapid Growth by Recognition of Peptidoglycan of Susceptible Competitors. Nat. Commun. 2022 131 2022, 13 (1), 1–15. 10.1038/s41467-021-27904-2.

(24) López, D.; Fischbach, M. A.; Chu, F.; Losick, R.; Kolter, R. Structurally Diverse Natural Products That Cause Potassium Leakage Trigger Multicellularity in Bacillus Subtilis. Proc. Natl. Acad. Sci. U. S. A. 2009, 106 (1), 280–285. 10.1073/PNAS.0810940106.

(25) Chin-A-Woeng, T. F. C.; Bloemberg, G. V.; Van Der Bij, A. J.; Van Der Drift, K. M. G. M.; Schripsema, J.; Kroon, B.; Scheffer, R. J.; Keel, C.; Bakker, P. A. H. M.; Tichy, H. V.; De Bruijn, F. J.; Thomas-Oates, J. E.; Lugtenberg, B. J. J. Biocontrol by Phenazine-1-Carboxamide-Producing Pseudomonas Chlororaphis PCL1391 of Tomato Root Rot Caused by Fusarium Oxysporum f. Sp. Radicis-Lycopersici. MPMI 1998, 11 (11), 1069–1077. 10.1094/MPMI.1998.11.11.1069.

(26) Dror, B.; Amutuhaire, H.; Frenkel, O.; Jurkevitch, E.; Cytryn, E. Identification of Bacterial Populations and Functional Mechanisms Potentially Involved in Biochar-Facilitated Antagonism of the Soilborne Pathogen Fusarium Oxysporum. Phytobiomes J. 2022, 6 (2), 139–150. 10.1094/PBIOMES-11-21-0071-R.

(27) Shanmugaiah, V.; Mathivanan, N.; Varghese, B. Purification, Crystal Structure and Antimicrobial Activity of Phenazine‐1‐carboxamide Produced by a Growth‐promoting Biocontrol Bacterium, Pseudomonas Aeruginosa MML2212. J. Appl. Microbiol. 2010, 108 (2), 703–711. 10.1111/J.1365-2672.2009.04466.X.

(28) Lyng, M.; Kovács, Á.T. Frenemies of the Soil: Bacillus and Pseudomonas Interspecies Interactions. Trends Microbiol. 2023, 31 (8), 845–857. 10.1016/J.TIM.2023.02.003.

(29) Molina-Santiago, C.; Pearson, J. R.; Navarro, Y.; Berlanga-Clavero, M. V.; Caraballo-Rodriguez, A. M.; Petras, D.; García-Martín, M. L.; Lamon, G.; Haberstein, B.; Cazorla, F. M.; de Vicente, A.; Loquet, A.; Dorrestein, P. C.; Romero, D. The Extracellular Matrix Protects Bacillus Subtilis Colonies from Pseudomonas Invasion and Modulates Plant Co-Colonization. Nat. Commun. 2019 101 2019, 10 (1), 1–15. 10.1038/s41467-019-09944-x.

(30) Kost, C.; Patil, K. R.; Friedman, J.; Garcia, S. L.; Ralser, M. Metabolic Exchanges Are Ubiquitous in Natural Microbial Communities. Nat. Microbiol. 2023 812 2023, 8 (12), 2244–2252. 10.1038/s41564-023-01511-x.

(31) Sun, X.; Xu, Z.; Xie, J.; Hesselberg-Thomsen, V.; Tan, T.; Zheng, D.; Strube, M. L.; Dragoš, A.; Shen, Q.; Zhang, R.; Kovács, Á.T. Bacillus Velezensis Stimulates Resident Rhizosphere Pseudomonas Stutzeri for Plant Health through Metabolic Interactions. ISME J. 2021 163 2021, 16 (3), 774–787. 10.1038/s41396-021-01125-3.

(32) Gingichashvili, S.; Duanis-Assaf, D.; Shemesh, M.; Featherstone, J. D. B.; Feuerstein, O.; Steinberg, D. The Adaptive Morphology of Bacillus Subtilis Biofilms: A Defense Mechanism against Bacterial Starvation. Microorg. 2020, Vol. 8, Page 62 2019, 8 (1), 62. 10.3390/MICROORGANISMS8010062.

(33) Pérez-Lorente, A. I.; Molina-Santiago, C.; de Vicente, A.; Romero, D. Sporulation Activated via σ W Protects Bacillus from a Tse1 Peptidoglycan Hydrolase Type VI Secretion System Effector. Microbiol. Spectr. 2023, 11 (2). 10.1128/spectrum.05045-22.

(34) Andrić, S.; Rigolet, A.; Argüelles Arias, A.; Steels, S.; Hoff, G.; Balleux, G.; Ongena, L.; Höfte, M.; Meyer, T.; Ongena, M. Plant-Associated Bacillus Mobilizes Its Secondary Metabolites upon Perception of the Siderophore Pyochelin Produced by a Pseudomonas Competitor. ISME J. 2022 172 2022, 17 (2), 263–275. 10.1038/s41396-022-01337-1.

(35) Chin-A-Woeng, T. F. C.; Thomas-Oates, J. E.; Lugtenberg, B. J. J.; Bloemberg, G. V. Introduction of the PhzH Gene of Pseudomonas Chlororaphis PCL1391 Extends the Range of Biocontrol Ability of Phenazine-1-Carboxylic Acid-Producing Pseudomonas Spp. Strains. MPMI 2001, 14 (8), 1006–1015. 10.1094/MPMI.2001.14.8.1006.

(36) Powers, M. J.; Sanabria-Valentín, E.; Bowers, A. A.; Shank, E. A. Inhibition of Cell Differentiation in Bacillus Subtilis by Pseudomonas Protegens. J. Bacteriol. 2015, 197 (13), 2129–2138. 10.1128/JB.02535-14.

(37) Andrić, S.; Meyer, T.; Rigolet, A.; Prigent-Combaret, C.; Höfte, M.; Balleux, G.; Steels, S.; Hoff, G.; De Mot, R.; McCann, A.; De Pauw, E.; Argüelles Arias, A.; Ongena, M. Lipopeptide Interplay Mediates Molecular Interactions between Soil Bacilli and Pseudomonads. Microbiol. Spectr. 2021, 9 (3). 10.1128/SPECTRUM.02038-21.

(38) Zhou, Y.; Wang, H.; Sun, J.; Wicaksono, W. A.; Liu, C.; He, Y.; Qin, Y.; Berg, G.; Li, L.; Lin, H.; Chai, Y.; Bai, Y.; Ma, Z.; Cernava, T.; Chen, Y. Phenazines Contribute to Microbiome Dynamics by Targeting Topoisomerase IV. Nat. Microbiol. 2025 1010 2025, 10 (10), 2396–2411. 10.1038/s41564-025-02118-0.

(39) Chen, Y.; Wang, J.; Yang, N.; Wen, Z.; Sun, X.; Chai, Y.; Ma, Z. Wheat Microbiome Bacteria Can Reduce Virulence of a Plant Pathogenic Fungus by Altering Histone Acetylation. Nat. Commun. 2018 91 2018, 9 (1), 3429-. 10.1038/s41467-018-05683-7.

(40) Pang, Z.; Chong, J.; Zhou, G.; De Lima Morais, D.A.; Chang, L.; Barrette, M.; Gauthier, C.; Jacques, P.É.; Li, S.; Xia, J. MetaboAnalyst 5.0: Narrowing the Gap between Raw Spectra and Functional Insights. Nucleic Acids Res. 2021, 49 (W1), W388–W396. 10.1093/NAR/GKAB382.

(41) Schmid, R.; Heuckeroth, S.; Korf, A.; Smirnov, A.; Myers, O.; Dyrlund, T. S.; Bushuiev, R.; Murray, K. J.; Hoffmann, N.; Lu, M.; Sarvepalli, A.; Zhang, Z.; Fleischauer, M.; Dührkop, K.; Wesner, M.; Hoogstra, S. J.; Rudt, E.; Mokshyna, O.; Brungs, C.; Ponomarov, K.; Mutabdžija, L.; Damiani, T.; Pudney, C. J.; Earll, M.; Helmer, P. O.; Fallon, T. R.; Schulze, T.; Rivas-Ubach, A.; Bilbao, A.; Richter, H.; Nothias, L. F.; Wang, M.; Orešič, M.; Weng, J. K.; Böcker, S.; Jeibmann, A.; Hayen, H.; Karst, U.; Dorrestein, P. C.; Petras, D.; Du, X.; Pluskal, T. Integrative Analysis of Multimodal Mass Spectrometry Data in MZmine 3. Nat. Biotechnol. 2023 414 2023, 41 (4), 447–449. 10.1038/s41587-023-01690-2.

(42) Wang, M.; Carver, J. J.; Phelan, V. V.; Sanchez, L. M.; Garg, N.; Peng, Y.; Nguyen, D. D.; Watrous, J.; Kapono, C. A.; Luzzatto-Knaan, T.; Porto, C.; Bouslimani, A.; Melnik, A. V.; Meehan, M. J.; Liu, W. T.; Crüsemann, M.; Boudreau, P. D.; Esquenazi, E.; Sandoval-Calderón, M.; Kersten, R. D.; Pace, L. A.; Quinn, R. A.; Duncan, K. R.; Hsu, C. C.; Floros, D. J.; Gavilan, R. G.; Kleigrewe, K.; Northen, T.; Dutton, R. J.; Parrot, D.; Carlson, E. E.; Aigle, B.; Michelsen, C. F.; Jelsbak, L.; Sohlenkamp, C.; Pevzner, P.; Edlund, A.; McLean, J.; Piel, J.; Murphy, B. T.; Gerwick, L.; Liaw, C. C.; Yang, Y. L.; Humpf, H. U.; Maansson, M.; Keyzers, R. A.; Sims, A. C.; Johnson, A. R.; Sidebottom, A. M.; Sedio, B. E.; Klitgaard, A.; Larson, C. B.; Boya, C. A. P.; Torres-Mendoza, D.; Gonzalez, D. J.; Silva, D. B.; Marques, L. M.; Demarque, D. P.; Pociute, E.; O’Neill, E. C.; Briand, E.; Helfrich, E. J. N.; Granatosky, E. A.; Glukhov, E.; Ryffel, F.; Houson, H.; Mohimani, H.; Kharbush, J. J.; Zeng, Y.; Vorholt, J. A.; Kurita, K. L.; Charusanti, P.; McPhail, K. L.; Nielsen, K. F.; Vuong, L.; Elfeki, M.; Traxler, M. F.; Engene, N.; Koyama, N.; Vining, O. B.; Baric, R.; Silva, R. R.; Mascuch, S. J.; Tomasi, S.; Jenkins, S.; Macherla, V.; Hoffman, T.; Agarwal, V.; Williams, P. G.; Dai, J.; Neupane, R.; Gurr, J.; Rodríguez, A. M. C.; Lamsa, A.; Zhang, C.; Dorrestein, K.; Duggan, B. M.; Almaliti, J.; Allard, P. M.; Phapale, P.; Nothias, L. F.; Alexandrov, T.; Litaudon, M.; Wolfender, J. L.; Kyle, J. E.; Metz, T. O.; Peryea, T.; Nguyen, D. T.; VanLeer, D.; Shinn, P.; Jadhav, A.; Müller, R.; Waters, K. M.; Shi, W.; Liu, X.; Zhang, L.; Knight, R.; Jensen, P. R.; Palsson, B.; Pogliano, K.; Linington, R. G.; Gutiérrez, M.; Lopes, N. P.; Gerwick, W. H.; Moore, B. S.; Dorrestein, P. C.; Bandeira, N. Sharing and Community Curation of Mass Spectrometry Data with Global Natural Products Social Molecular Networking. Nat. Biotechnol. 2016 348 2016, 34 (8), 828–837. 10.1038/nbt.3597.

(43) Dührkop, K.; Fleischauer, M.; Ludwig, M.; Aksenov, A. A.; Melnik, A. V.; Meusel, M.; Dorrestein, P. C.; Rousu, J.; Böcker, S. SIRIUS 4: A Rapid Tool for Turning Tandem Mass Spectra into Metabolite Structure Information. Nat. Methods 2019 164 2019, 16 (4), 299–302. 10.1038/s41592-019-0344-8.

(44) Dunn, W. B.; Erban, A.; Weber, R. J. M.; Creek, D. J.; Brown, M.; Breitling, R.; Hankemeier, T.; Goodacre, R.; Neumann, S.; Kopka, J.; Viant, M. R. Mass Appeal: Metabolite Identification in Mass Spectrometry-Focused Untargeted Metabolomics. Metabolomics 2013, 9 (SUPPL.1), 44–66. 10.1007/s11306-012-0434-4.

(45) Carvalho, M. P. de; Abraham, W.-R. Antimicrobial and Biofilm Inhibiting Diketopiperazines. Curr. Med. Chem. 2012, 19 (21), 3564–3577. 10.2174/092986712801323243.

(46) Wang, G.; Dai, S.; Chen, M.; Wu, H.; Xie, L.; Luo, X.; Li, X. Two Diketopiperazine Cyclo(PRO-PHE) Isomers from Marine Bacteria Bacillus Subtilis Sp. 13-2. Chem. Nat. Compd. 2010, 46 (4), 583–585. 10.1007/S10600-010-9680-8.

(47) Elkahoui, S.; Rahim, H. A.; Tabbene, O.; Shaaban, M.; Limam, F.; Laatsch, H. Cyclo-(His,Leu): A New Microbial Diketopiperazine from a Terrestrial Bacillus Subtilis Strain B38. Nat. Prod. Res. 2013, 27 (2), 108–116. 10.1080/14786419.2012.660635.

(48) Albarracín Orio, A.G.; Petras, D.; Tobares, R. A.; Aksenov, A. A.; Wang, M.; Juncosa, F.; Sayago, P.; Moyano, A. J.; Dorrestein, P. C.; Smania, A. M. Fungal–Bacterial Interaction Selects for Quorum Sensing Mutants with Increased Production of Natural Antifungal Compounds. Commun. Biol. 2020 31 2020, 3 (1), 1–9. 10.1038/s42003-020-01342-0.

(49) Albarracín Orio, A.G.; Brücher, E.; Ducasse, D. A. A Strain of Bacillus Subtilis Subsp. Subtilis Shows a Specific Antagonistic Activity against the Soil-Borne Pathogen of Onion Setophoma Terrestris. Eur. J. Plant Pathol. 2016, 144 (1), 217–223. 10.1007/s10658-015-0762-0.

(50) Mavrodi, D. V.; Mavrodi, O. V.; Parejko, J. A.; Bonsall, R. F.; Kwak, Y. S.; Paulitz, T. C.; Thomashow, L. S.; Weller, D. M. Accumulation of the Antibiotic Phenazine-1-Carboxylic Acid in the Rhizosphere of Dryland Cereals. Appl. Environ. Microbiol. 2012, 78 (3), 804–812. 10.1128/AEM.06784-11.

(51) Zhao, X.; Drlica, K. Reactive Oxygen Species and the Bacterial Response to Lethal Stress. Curr. Opin. Microbiol. 2014, 21, 1–6. 10.1016/J.MIB.2014.06.008.

(52) Borisov, V. B.; Siletsky, S. A.; Nastasi, M. R.; Forte, E. ROS Defense Systems and Terminal Oxidases in Bacteria. Antioxidants 2021, Vol. 10, Page 839 2021, 10 (6), 839. 10.3390/ANTIOX10060839.

(53) Zhang, L.; Sun, C. Fengycins, Cyclic Lipopeptides from Marine Bacillus Subtilis Strains, Kill the Plant-Pathogenic Fungus Magnaporthe Grisea by Inducing Reactive Oxygen Species Production and Chromatin Condensation. Appl. Environ. Microbiol. 2018, 84 (18), 445–463. 10.1128/AEM.00445-18.

(54) Tang, Q.; Bie, X.; Lu, Z.; Lv, F.; Tao, Y.; Qu, X. Effects of Fengycin from Bacillus Subtilis FmbJ on Apoptosis and Necrosis in Rhizopus Stolonifer. J. Microbiol. 2014, 52 (8), 675–680. 10.1007/S12275-014-3605-3.

(55) Lee, J. W.; Helmann, J. D. The PerR Transcription Factor Senses H2O2 by Metal-Catalysed Histidine Oxidation. Nat. 2006 4407082 2006, 440 (7082), 363–367. 10.1038/nature04537.

