## Supporting information including experimental procedures and figures S1-S12 for "Chemical Crosstalk between the Common Soil Bacteria *Pseudomonas chlororaphis* and *Bacillus subtilis*"

#### Bacterial growth media and conditions

*P. chlororaphis* PCL 1391 (*P. chlororaphis* WT) and *P. chlororaphis* PCL 1119 (*P. chlororaphis* Δ*phz*) were kindly provided to us by Dianne K. Newman (California Institute of Technology, Pasadena, California, USA). *B. subtilis* NCIB 3610 (*B. subtilis* WT), *B. subtilis* Δ*pps* and *B. subtilis* Δ*srfAA* were provided by Ilana Kolodkin-Gal (Reichman University, Herzliya, Israel), and *F. oxysporum* strain f. sp. *radicis-cucumerinum* was provided by Omer Frenkel (Institute of Plant Protection, Agricultural Research Organization (ARO), The Volcani Center). All strains were grown on LB media at 30˚C. An overnight liquid culture (30˚C, 200 RPM) was diluted to $OD_{600}$=0.5 for the solid cultures experiment, and 2 µL was plated on slim LB agar plates. For interaction assays, the bacteria were plated at a distance of 0.8 cm, unless specified otherwise, and incubated at 30˚C for 48 hr. PCN was added to the media to achieve final concentrations of 10 µM to the required plates before the solidification of the agar (same concentration was maintained in all plate setups, except specified otherwise). For colony forming unit (CFU) experiments, the bacteria were diluted in PBS solution and plated on isolation media, nutrient agar with polymyxin B, and *Pseudomonas* isolation agar for *B. subtilis* and *P. chlororaphis*, respectively. The colonies were counted after 24 hours.

#### Metabolites extraction for metabolomics analysis

Four replicates of the bacteria colonies were cut from the plates with the agar in the same size and incubated for 1 hr at room temperature in 1 mL LC\MS grade MeOH (Baker Analyzed® LC-MS reagent, J.T. Baker). Followed by centrifuge for 10 min in 9400 RFC (Eppendorf™ 5424 Microcentrifuges). The supernatant was mixed with LC\MS grade water (Baker Analyzed® LC-MS reagent, J.T. Baker) in 1:1 ratio, filtered with 0.22 µm PVDF filter and was kept in -20˚C until further analysis. The procedure was done for three different biological repeats.

#### PCN calibration curve preparation

Phenazine-1-carboxamide (PCN) was dissolved in MeOH:water 1:1 (Baker Analyzed® LC-MS reagent, J.T. Baker), and then diluted to the required concentrations. PCN concentration was measured in technical triplicates, and the average was used to create the calibration curve.

#### Liquid chromatography-tandem mass spectrometry (LC-MS/MS)

Samples were examined by liquid chromatography-tandem mass spectrometry (LC-MS/MS) using an UltiMate 3000 UHPLC+ focused LC-MS system (Thermo Scientific™) occupied with Accucore™ C18 HPLC Columns (2.6 µm particle size, 2.1 mmx 100 mm) (Thermo Scientific™), followed by Q Exactive™ Focus Hybrid Quadrupole-Orbitrap™ Mass Spectrometer (Thermo Scientific™) with HESI ion source. The flow rate was set to 0.4 mL/min when solvent A was water (Baker Analyzed® LC-MS reagent, J.T. Baker) with 0.1% formic acid (99% ULC/MS - CC/SFC, Bio-Lab), and solvent B was acetonitrile (Baker Analyzed® LC-MS reagent, J.T. Baker) with 0.1% formic acid. 5 µL was injected to the preheated column oven to 40˚C. Gradients started with 100% of solvent A for 2 min. After 2 min the gradient was increased to 50% solvent B, and in the following 6 min the gradient was increased to 90% solvent B and stayed in this composition for 1 min. The gradient changed to 20% solvent B over the next minute, and was maintained that way for another minute. The scan range was set to 100-1500 m/z, with a normalized collision energy of 30 eV. The capillary temperature was 300˚C, and the spray voltage was 3.8 kV. To evaluate the instrument operations, the four first and last samples were a quality control (QC) sample (composed of all the samples). QC sample was tested every 10 experimental samples. MS spectra were analyzed by Xcalibur and freestyle softwares (Thermo Scientific™) when analyzed manually.

**Mass spectrometry data analysis**

Acquired MS .raw files were converted to .mzML files with proteowizard (MSConvert).[1] The converted files were uploaded to MZmine3[2] for feature detection and alignment. MZmine3 output was uploaded to GNPS[3] and Sirius[4] for further analysis. To learn more about the relationships and structures of the metabolites in the cultures, molecular networks were created in GNPS[3,5] with precursor ion mass tolerance and fragment ion mass tolerance of 0.02 Da, and cosine score of 0.7 to form a network edge. The Sirius tool,[6–9] with CSI:FingerID and CANOPUS, was used to annotate a putative identification of the compounds' structure. MetaboAnalyst[10] was used to create a principal component analysis based on the $MS^1$ data.[#]

**Plipastatin and surfactin extraction**

An overnight culture of *B. subtilis* WT was plated on slim LB plates. After two days, the plates were inserted into centrifuge tubes with MeOH, vortexed, and kept at room temperature for 1 hr. The tubes were centrifuged (4˚C, 3200 g, 20 min) and concentrated in a vacuum. Before the solvent evaporated completely, it was filtered via a 0.22 µm PVDF filter. The resulting pellet was extracted using solid phase extraction (SPE) method by the manufacture protocol using Biotage® VacMaster™ processing manifold. Briefly, SPE columns (Strata™-X 33µm Polymeric Reversed Phase 30mg, phenomenex®) were conditioned and equilibrated with 1 mL MeOH and 1 mL water, respectively. Then, the supernatant was loaded onto the columns; the columns were washed with 1 mL water and dried. The metabolites were eluted from the column with 500 µL 2% formic acid in MeOH twice. The samples were concentrated in a vacuum (CentriVap Concentrators, LABCONCO) and kept at -20˚C until purification. The compounds were separated using UltiMate 3000 HPLC (Thermo Scientific™) occupied with Luna® C18(2) 100 Å LC Column (250 X 21.2 mm, 5 µm particle size) (phenomenex®). The flow rate was set to 20 mL/min when solvent A was water (Milli-Q®, Millipore) with 0.1% trifluoracetic acid (ULC/MS - CC/SFC, Bio-Lab), and solvent B was acetonitrile (HPLC, J.T. Baker) with 0.1% trifluoracetic acid. The gradient started with 100% of solvent A for 3 min. After 6 min, the gradient was increased to 50% solvent A, and in the following 15 min, the gradient was increased to 90% solvent A and stayed in this composition for 5 min. The gradient changed to 20% over the next minute and was maintained for another 3 min. The UV channel was set to 205 nm. The composition of each fraction was assessed by LC-MS as described above, and concentrations were measured using a nanodrop spectrophotometer (NanoDrop™ 2000c, Thermo Scientific™).

**Metabolite tolerance assay**

An overnight culture of *B. subtilis* WT, *B. subtilis* Δ*pps*, and *B. subtilis* Δ*srfAA* was diluted to $OD_{600}$=0.1 and added to a serial dilution of PCN in 96 well plate (Corning™ Costar™). Similarly, *P. chlororaphis* WT was introduced to a purified plipastatin derivate or surfactin mixture. The plate was sealed with a breathable sealing membrane (Breathe-Easy®, Merk). $OD_{600}$ was measured every 20 min for 13-16 hr in a plate reader (Varioskan™ LUX, Thermo Scientific™), while the plate was shaken between measurements.

#### Reactive oxygen species assay

An overnight starter of *P. chlororaphis* WT was diluted and incubated for 6 hr at 30°C with shaking. After 6 hr, the bacteria were diluted 1:10 with fresh LB, and 180 µL was transferred to each well of a black 96 well plate (chimney well, Greiner bio-one, Germany). 10 µL from the desired compounds were added to the wells, and water was added to the control wells. Finally, 10 µL of 2′,7′-dichlorodihydrofluorescein diacetate (DCFDA) 1 mM was added to each well. DCFDA is a cell-permeable compound. Inside the cell, it is cleaved by cellular esterases. In the presence of reactive oxygen species (ROS), the cleavaged compound oxidized to the fluorescent compound 2',7'-dichlorofluorescein (DCF). $OD_{600}$ and relative fluorescence were measured by a plate reader (Varioskan™ LUX, Thermo Scientific™). The samples were excited at 485 nm, and emission was measured at 535 nm. The relative fluorescence was divided by the $OD_{600}$ of each well.

#### Effect of *P. chlororaphis* and *B. subtilis* interaction on the growth of *Fusarium oxysporum*

#### Microbial strains and culture conditions

*P. chlororaphis* WT and *B. subtilis* WT were cultured individually and in combination (coculture) in Luria–Bertani (LB) broth at 30 °C under constant agitation (150 rpm) for 48 hours. Cultures were standardized to an optical density of $OD_{600} = 0.2$ before use. Following incubation, the cultures were centrifuged at 10,000 × g for 10 minutes, and the supernatants were filter-sterilized using 0.22 µm membrane filters (Millipore) to obtain cell-free culture supernatants (CFS). *Fusarium oxysporum* was grown on Potato Dextrose Agar (PDA) and incubated at 28 ± 2 °C for 7 days prior to the experiment. CFU was conducted as describe above.

#### Antagonism assay using bacterial supernatants

To evaluate the effects of bacterial metabolites on fungal growth, a modified in situ PDA-CFS assay was employed[11]. Melted 6% PDA was cooled to 45 °C and mixed at a 1:1 (v/v) ratio with either axenic bacterial or coculture CFS. The resulting mixtures were poured into sterile Petri dishes and allowed to solidify under aseptic conditions. A 6 mm diameter agar plug was aseptically excised from the actively growing edge of a 7-day-old *F. oxysporum* colony and placed at the center of each Petri dish. Control plates were prepared using PDA mixed with sterile LB medium processed identically to the treatments. Plates were incubated at 28 ± 2 °C, and fungal growth was monitored for 6 days (144 hours). Fungal colony diameter was measured from Day 3 onward using a transparent ruler and the ImageJ web-based platform ([https://ij.imjoy.io](https://ij.imjoy.io)) for inhibition analysis. To assess the bacterial CFS effect on the fungal growth dynamics, the radial growth rate (Kr) was measured and the inhibition percentage relative to the untreated control was calculated[12].

$$Kr = \frac{r2-r1}{t2-t1} \quad \text{.....eq1}$$

$$\%\, inhibition = \left(\frac{control-treatment}{control}\right) x\, 100 \text{ ....................................... eq2}$$

Where **r** is the average colony radius (mm) at time points ***t1*** and ***t2***. All treatments were performed in triplicate. The mean values were used for graphical presentation and statistical analysis.

**Supporting figures**

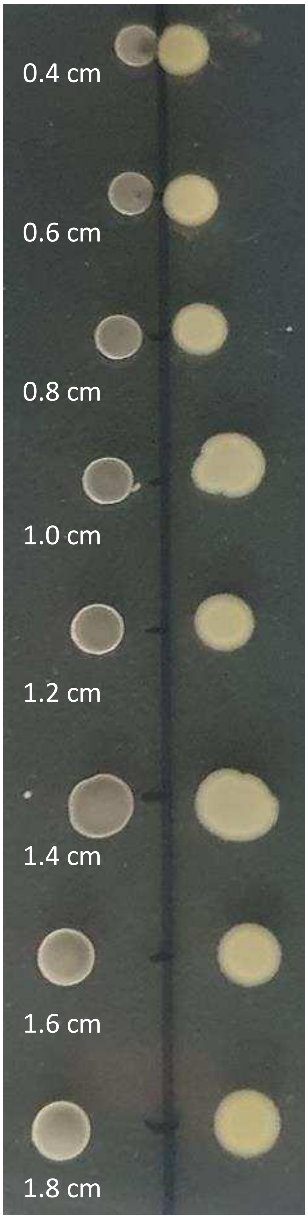


**Figure S1.** *B. subtilis* WT and *P. chlororaphis* WT co-cultures. Below each pair the initial plating distance is noted.

**A**

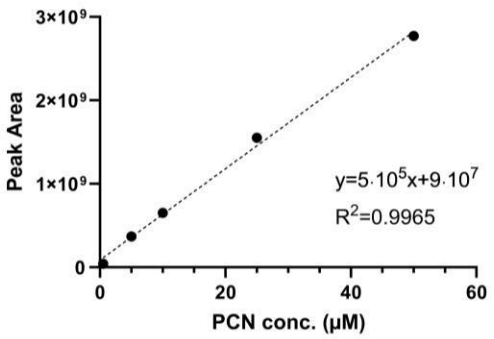


**B**

| Peak area | PCN conc. (μM) | Average (μM) | Standard deviation (μM) |
|---|---|---|---|
| 6.47E+08 | 11.14 | 11.45 | 6.62 |
| 4.95E+08 | 8.11 | | |
| 1.13E+09 | 20.81 | | |
| 3.77E+08 | 5.74 | | |

**Figure S2.** PCN production by *P. chlororaphis* on solid media. (A) calibration carve (B) measured PCN quantities.

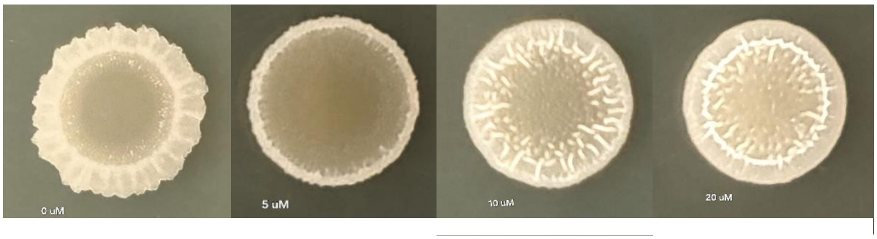


0 µM 5 µM 10 µM 20 µM

**Figure S3.** The effect of PCN at different concentrations on *B. subtills* morphology.

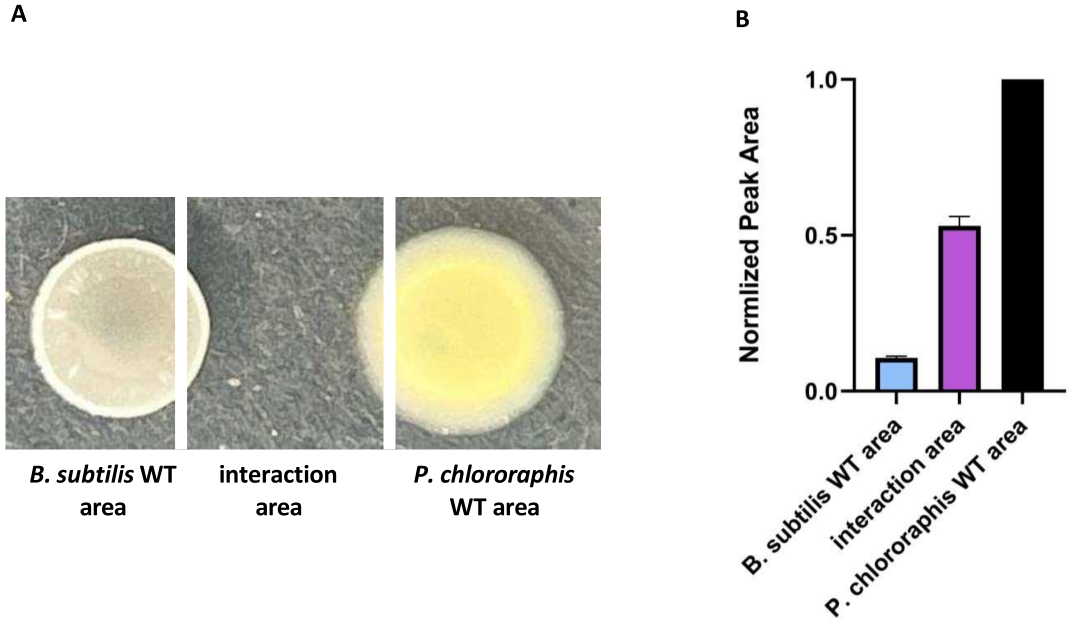


**Figure S4.** PCN concentration by culture area. (A) the divided areas (B) Peak area of PCN. The data represents the average of normalized peak area of three biological replicates, consist of four technical replicates, normalized base on PCN concentrations at the *P. chlororaphis* WT area.

To further explore whether the PCN secreted by *P. chlororaphis* causes the visible change in *B. subtilis* morphology by spreading to the whole colony, or whether the signal is detected by *B. subtilis* only near the interaction zone, we cut the colonies into three parts and analyzed PCN concentrations in each zone separately: (i) the first consists only of *B. subtilis*, (ii) the second zone is the interaction area between the bacteria, including parts of both the colonies, (iii) the third zone consists only of *P. chlororaphis* (Figure S3). We detected the highest concentrations of PCN in the *P. chlororaphis* section, while the lowest concentration was in the *B. subtilis* sections. As we observed no indication of a transformation of PCN by *B. subtilis*, we assume that even a low concentration of PCN is detected by *B. subtilis*, causing these morphological changes. Whether the relatively small concentrations of PCN reaching the whole *B. subtilis* colony are enough to affect the morphology of the entire colony is not clear. It is possible that the higher concentrations detected in the interaction area between the bacteria can cause a signaling cascade that eventually reaches the entire *B. subtilis* colony. Further investigations are needed in order to examine the precise mode of signal transduction.

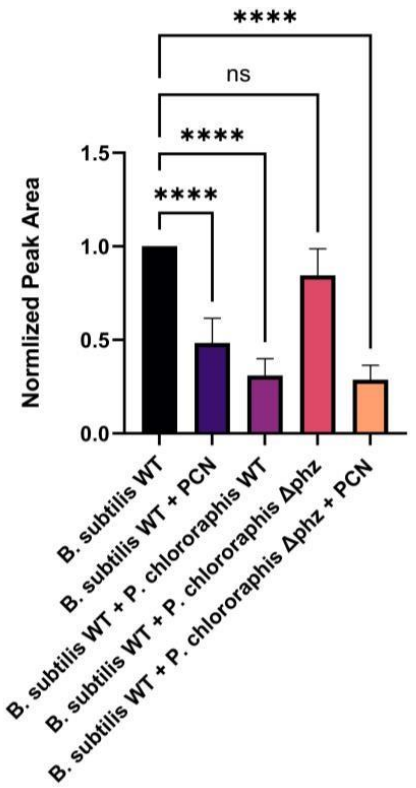


**Figure S5.** Production of a compound with m/z of 227.1754 by *B. subtilis* WT in various treatments. The data represent the average of normalized peak area of three biological replicates, consisting of four technical replicates, based on the compound production in mono-cultures of *B. subtilis* WT. Statistical analysis was performed using one-way ANOVA, with $p<0.0001$ marked by ****.

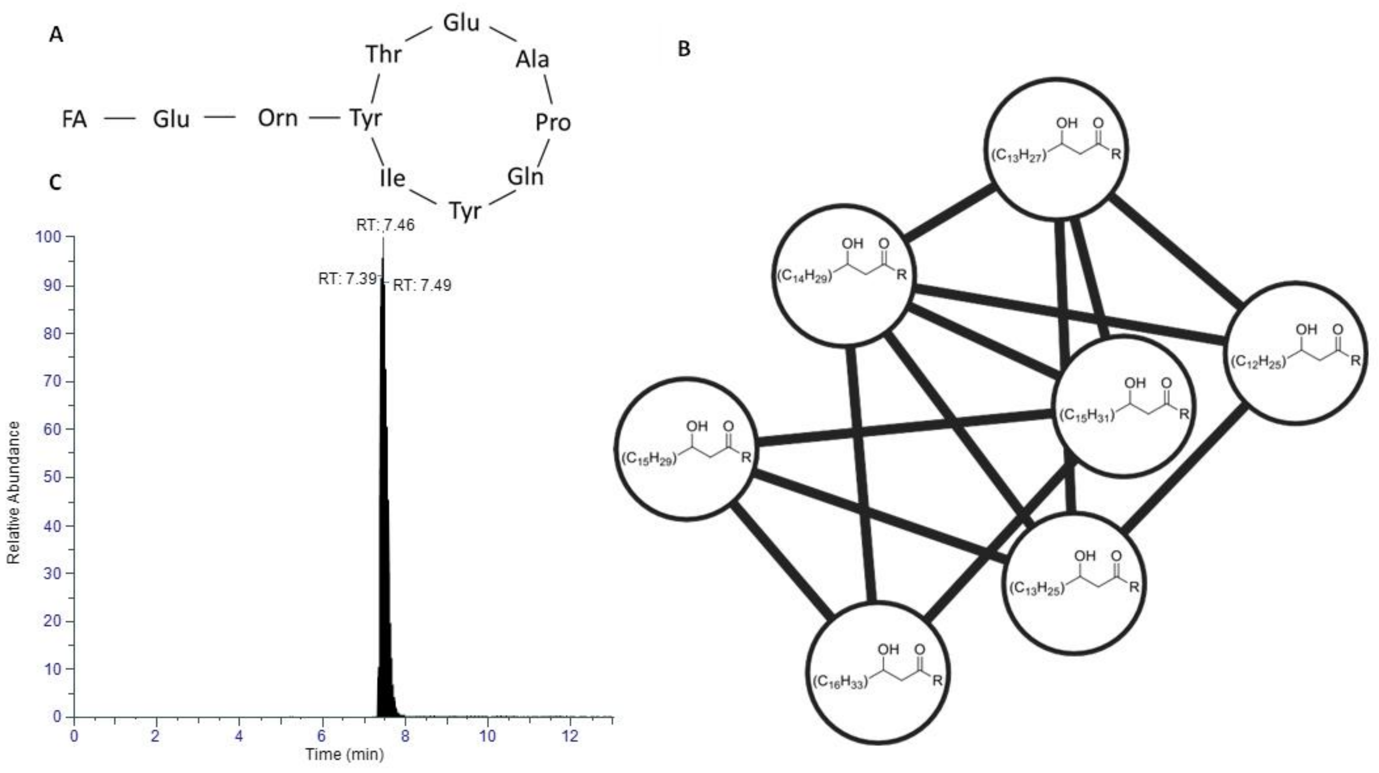


**Figure S6.** Plipastatin derivative A $C_{19}$ extraction. (A) Plipastatin basic structure. (B) Molecular networking cluster for plipastatin. The fatty acids (FA) suitable for the detected masses are shown inside the appropriate node. Each node represents a metabolite in the sample, and any edge connecting two nodes suggests that they have a similar structure. (C) Chromatograms of the extracted plipastatin.

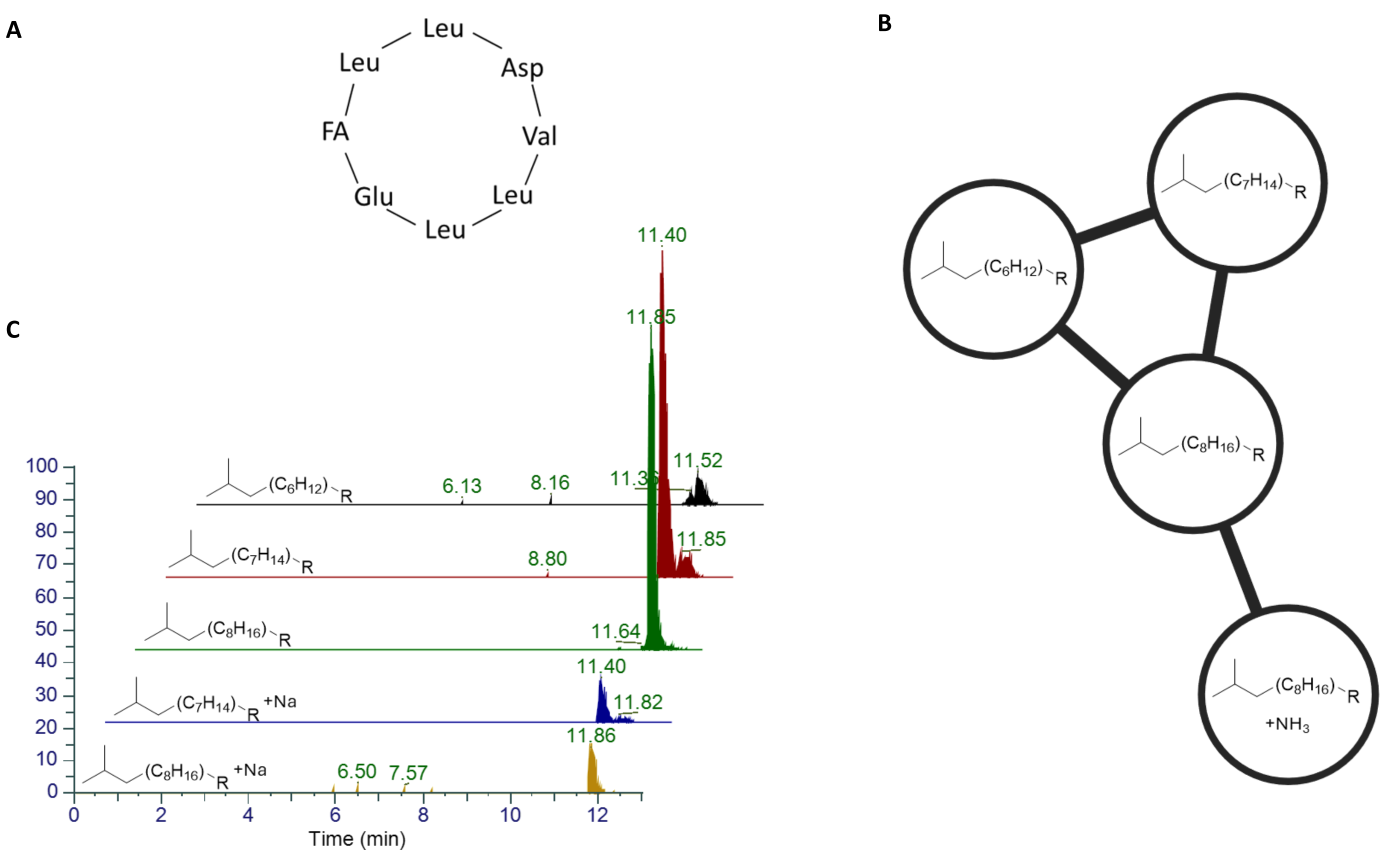


**Figure S7.** Surfactin extraction. (A) Surfactin basic structure. (B) Molecular networking cluster for surfactin. The fatty acids (FA) suitable for the detected masses are shown inside the appropriate node. Each node represents a metabolite in the sample, and any edge connecting two nodes suggests that they have a similar structure. (C) Chromatograms of the extracted surfactins, the suitable FA it presented next to the chromatograms. (B-C) only adducts different from H are mentioned.

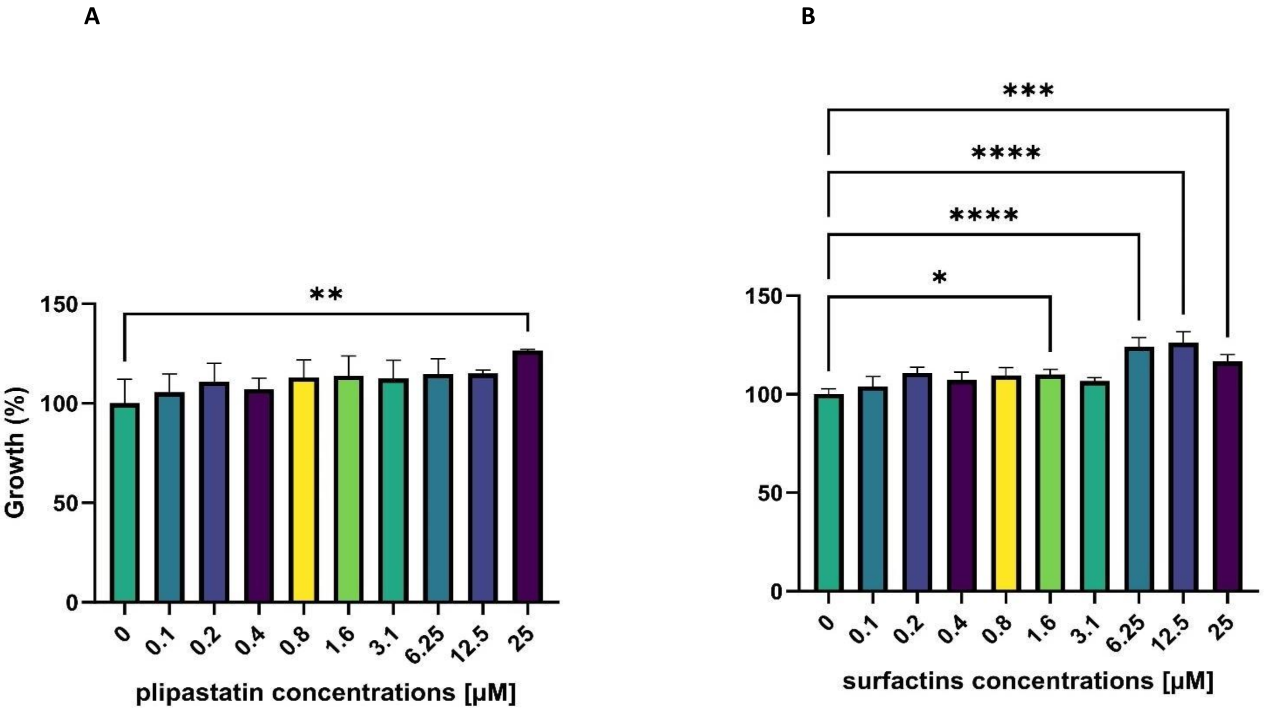


**Figure S8.** *P. chlororaphis* WT growth in presence of (A) plipastatin derivative and (B) different surfactins mixture concentration at 70% from maximum growth. Error bars represent standard deviation of triplicates. Statistical analysis was performed using one-way ANOVA, with p<0.0001, p<0.001, P<0.01 and p<0.05 marked by ****, ***, **, *, respectively.

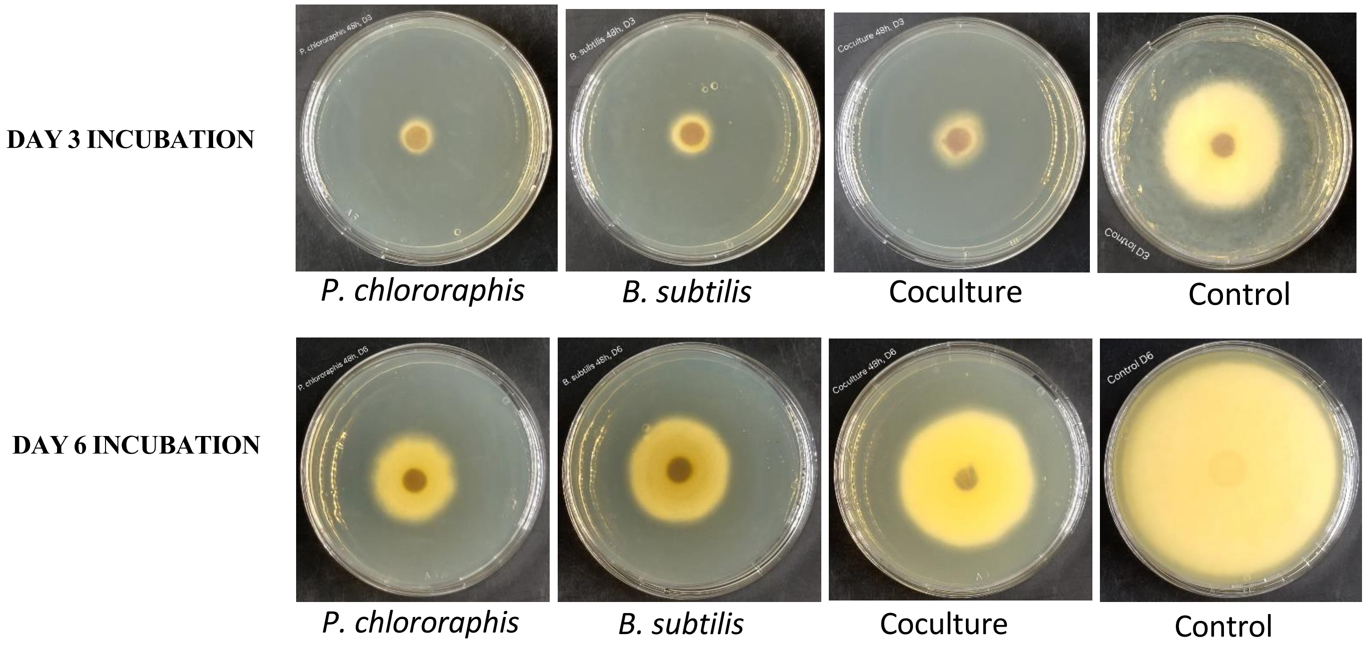


**Figure S9**. Growth of fungal colonies on agar that contains supernatants of mature bacterial cultures. Visual inspection of the colonies showed inhibition in the presence of all bacterial supernatants (compared to controls), but enhanced growth of *F. oxysporum* in the coculture CFS compared to the individual bacterial treatments.

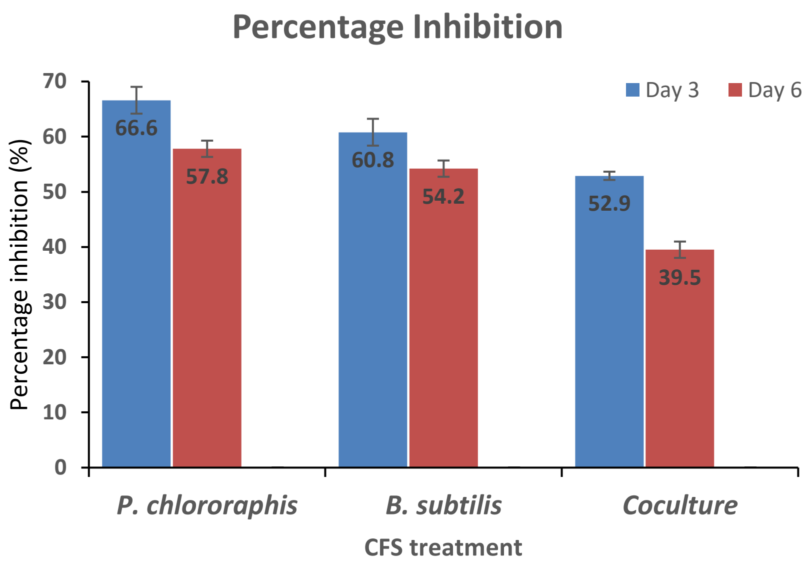


**Figure S10**. On both day 3 and day 6, the coculture supernatant exhibited the least inhibitory effect, while *P. chlororaphis* CFS showed the most inhibition, followed by *B. subtilis*. Values represent the mean of three setups and error bars represent standard deviation of the triplicates.

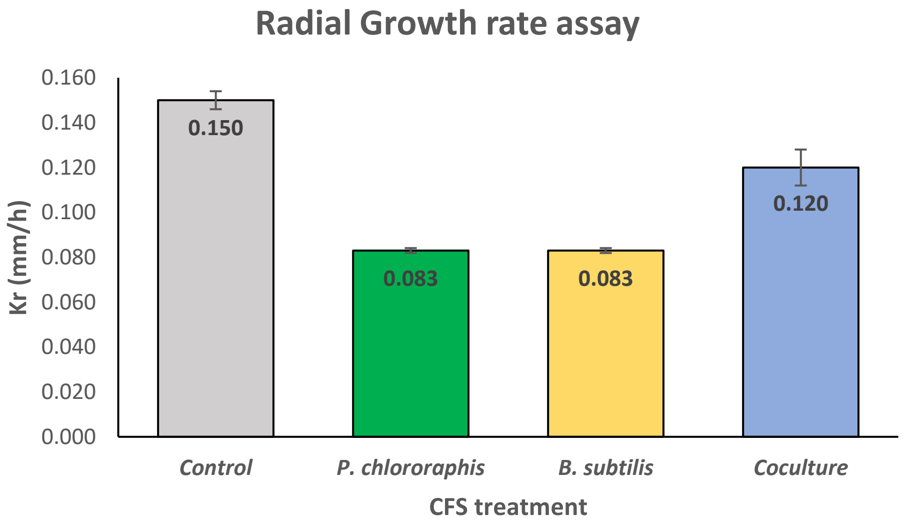


**Figure S11.** Radial growth rate (Kr) analysis shows that the control fungus exhibited nearly twice the radial growth rate compared to those grown on media containing individual bacterial CFS. The coculture treatment resulted in a higher Kr value than either individual treatment, indicating a potential conservation of inhibitory compound production by *P. chlororaphis* and *B. subtilis* that reduced their combined antifungal efficacy. Values represent the mean of three setups, and error bars represent the standard deviation of the triplicates.

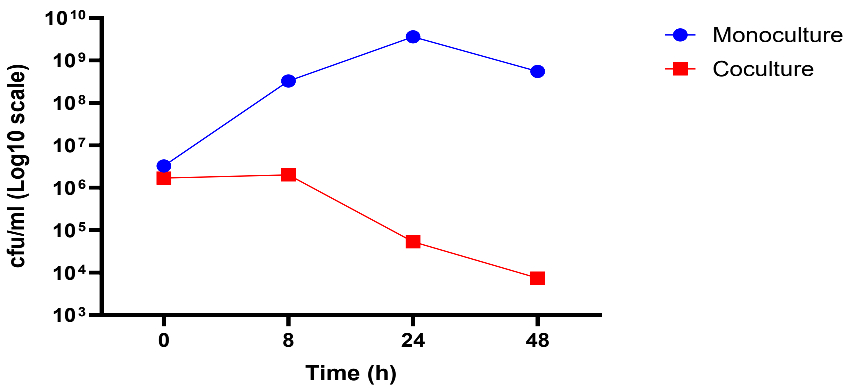


**Figure S12.** *B. subtilis* viability in liquid LB cultures in mono-culture vs. co-culture with *P. chlororaphis*. This indicates that *B. subtilis* can survive for up to 48 hours in liquid media with *P. chlororaphis*, despite a decrease in population. Values represent the mean of three setups and error bars represent standard deviation of the triplicates.

### References

(1) Chambers, M. C.; MacLean, B.; Burke, R.; Amodei, D.; Ruderman, D. L.; Neumann, S.; Gatto, L.; Fischer, B.; Pratt, B.; Egertson, J.; Hoff, K.; Kessner, D.; Tasman, N.; Shulman, N.; Frewen, B.; Baker, T. A.; Brusniak, M. Y.; Paulse, C.; Creasy, D.; Flashner, L.; Kani, K.; Moulding, C.; Seymour, S. L.; Nuwaysir, L. M.; Lefebvre, B.; Kuhlmann, F.; Roark, J.; Rainer, P.; Detlev, S.; Hemenway, T.; Huhmer, A.; Langridge, J.; Connolly, B.; Chadick, T.; Holly, K.; Eckels, J.; Deutsch, E. W.; Moritz, R. L.; Katz, J. E.; Agus, D. B.; MacCoss, M.; Tabb, D. L.; Mallick, P. A Cross-Platform Toolkit for Mass Spectrometry and Proteomics. *Nat. Biotechnol. 2012 3010* **2012**, *30* (10), 918–920. https://doi.org/10.1038/nbt.2377.

(2) Schmid, R.; Heuckeroth, S.; Korf, A.; Smirnov, A.; Myers, O.; Dyrlund, T. S.; Bushuiev, R.; Murray, K. J.; Hoffmann, N.; Lu, M.; Sarvepalli, A.; Zhang, Z.; Fleischauer, M.; Dührkop, K.; Wesner, M.; Hoogstra, S. J.; Rudt, E.; Mokshyna, O.; Brungs, C.; Ponomarov, K.; Mutabdžija, L.; Damiani, T.; Pudney, C. J.; Earll, M.; Helmer, P. O.; Fallon, T. R.; Schulze, T.; Rivas-Ubach, A.; Bilbao, A.; Richter, H.; Nothias, L. F.; Wang, M.; Orešič, M.; Weng, J. K.; Böcker, S.; Jeibmann, A.; Hayen, H.; Karst, U.; Dorrestein, P. C.; Petras, D.; Du, X.; Pluskal, T. Integrative Analysis of Multimodal Mass Spectrometry Data in MZmine 3. *Nat. Biotechnol. 2023 414* **2023**, *41* (4), 447–449. https://doi.org/10.1038/s41587-023-01690-2.

(3) Wang, M.; Carver, J. J.; Phelan, V. V.; Sanchez, L. M.; Garg, N.; Peng, Y.; Nguyen, D. D.; Watrous, J.; Kapono, C. A.; Luzzatto-Knaan, T.; Porto, C.; Bouslimani, A.; Melnik, A. V.; Meehan, M. J.; Liu, W. T.; Crüsemann, M.; Boudreau, P. D.; Esquenazi, E.; Sandoval-Calderón, M.; Kersten, R. D.; Pace, L. A.; Quinn, R. A.; Duncan, K. R.; Hsu, C. C.; Floros, D. J.; Gavilan, R. G.; Kleigrewe, K.; Northen, T.; Dutton, R. J.; Parrot, D.; Carlson, E. E.; Aigle, B.; Michelsen, C. F.; Jelsbak, L.; Sohlenkamp, C.; Pevzner, P.; Edlund, A.; McLean, J.; Piel, J.; Murphy, B. T.; Gerwick, L.; Liaw, C. C.; Yang, Y. L.; Humpf, H. U.; Maansson, M.; Keyzers, R. A.; Sims, A. C.; Johnson, A. R.; Sidebottom, A. M.; Sedio, B. E.; Klitgaard, A.; Larson, C. B.; Boya, C. A. P.; Torres-Mendoza, D.; Gonzalez, D. J.; Silva, D. B.; Marques, L. M.; Demarque, D. P.; Pociute, E.; O'Neill, E. C.; Briand, E.; Helfrich, E. J. N.; Granatosky, E. A.; Glukhov, E.; Ryffel, F.; Houson, H.; Mohimani, H.; Kharbush, J. J.; Zeng, Y.; Vorholt, J. A.; Kurita, K. L.; Charusanti, P.;

McPhail, K. L.; Nielsen, K. F.; Vuong, L.; Elfeki, M.; Traxler, M. F.; Engene, N.; Koyama, N.; Vining, O. B.; Baric, R.; Silva, R. R.; Mascuch, S. J.; Tomasi, S.; Jenkins, S.; Macherla, V.; Hoffman, T.; Agarwal, V.; Williams, P. G.; Dai, J.; Neupane, R.; Gurr, J.; Rodríguez, A. M. C.; Lamsa, A.; Zhang, C.; Dorrestein, K.; Duggan, B. M.; Almaliti, J.; Allard, P. M.; Phapale, P.; Nothias, L. F.; Alexandrov, T.; Litaudon, M.; Wolfender, J. L.; Kyle, J. E.; Metz, T. O.; Peryea, T.; Nguyen, D. T.; VanLeer, D.; Shinn, P.; Jadhav, A.; Müller, R.; Waters, K. M.; Shi, W.; Liu, X.; Zhang, L.; Knight, R.; Jensen, P. R.; Palsson, B.; Pogliano, K.; Linington, R. G.; Gutiérrez, M.; Lopes, N. P.; Gerwick, W. H.; Moore, B. S.; Dorrestein, P. C.; Bandeira, N. Sharing and Community Curation of Mass Spectrometry Data with Global Natural Products Social Molecular Networking. *Nat. Biotechnol. 2016 348* **2016**, *34* (8), 828–837. https://doi.org/10.1038/nbt.3597.

(4) Dührkop, K.; Fleischauer, M.; Ludwig, M.; Aksenov, A. A.; Melnik, A. V.; Meusel, M.; Dorrestein, P. C.; Rousu, J.; Böcker, S. SIRIUS 4: A Rapid Tool for Turning Tandem Mass Spectra into Metabolite Structure Information. *Nat. Methods 2019 164* **2019**, *16* (4), 299–302. https://doi.org/10.1038/s41592-019-0344-8.

(5) Schmid, R.; Petras, D.; Nothias, L. F.; Wang, M.; Aron, A. T.; Jagels, A.; Tsugawa, H.; Rainer, J.; Garcia-Aloy, M.; Dührkop, K.; Korf, A.; Pluskal, T.; Kameník, Z.; Jarmusch, A. K.; Caraballo-Rodríguez, A. M.; Weldon, K. C.; Nothias-Esposito, M.; Aksenov, A. A.; Bauermeister, A.; Albarracin Orio, A.; Grundmann, C. O.; Vargas, F.; Koester, I.; Gauglitz, J. M.; Gentry, E. C.; Hövelmann, Y.; Kalinina, S. A.; Pendergraft, M. A.; Panitchpakdi, M.; Tehan, R.; Le Gouellec, A.; Aleti, G.; Mannochio Russo, H.; Arndt, B.; Hübner, F.; Hayen, H.; Zhi, H.; Raffatellu, M.; Prather, K. A.; Aluwihare, L. I.; Böcker, S.; McPhail, K. L.; Humpf, H. U.; Karst, U.; Dorrestein, P. C. Ion Identity Molecular Networking for Mass Spectrometry-Based Metabolomics in the GNPS Environment. *Nat. Commun. 2021 121* **2021**, *12* (1), 1–12. https://doi.org/10.1038/s41467-021-23953-9.

(6) Sansinenea, E. *Bacillus* Spp.: As Plant Growth-Promoting Bacteria. *Second. Metab. Plant Growth Promot. Rhizomicroorganisms Discov. Appl.* **2019**, 225–237. https://doi.org/10.1007/978-981-13-5862-3_11.

(7) Djoumbou Feunang, Y.; Eisner, R.; Knox, C.; Chepelev, L.; Hastings, J.; Owen, G.; Fahy, E.; Steinbeck, C.; Subramanian, S.; Bolton, E.; Greiner, R.; Wishart, D. S. ClassyFire: Automated Chemical Classification with a Comprehensive, Computable Taxonomy. *J. Cheminform.* **2016**, *8* (1), 1–20. https://doi.org/10.1186/s13321-016-0174-y.

(8) Dührkop, K.; Shen, H.; Meusel, M.; Rousu, J.; Böcker, S. Searching Molecular Structure Databases with Tandem Mass Spectra Using CSI:FingerID. *Proc. Natl. Acad. Sci. U. S. A.* **2015**, *112* (41), 12580–12585. https://doi.org/10.1073/pnas.1509788112.

(9) Kim, H. W.; Wang, M.; Leber, C. A.; Nothias, L. F.; Reher, R.; Kang, K. Bin; Van Der Hooft, J. J. J.; Dorrestein, P. C.; Gerwick, W. H.; Cottrell, G. W. NPClassifier: A Deep Neural Network-Based Structural Classification Tool for Natural Products. *J. Nat. Prod.* **2021**, *84* (11), 2795–2807. https://doi.org/10.1021/acs.jnatprod.1c00399.

(10) Pang, Z.; Zhou, G.; Ewald, J.; Chang, L.; Hacariz, O.; Basu, N.; Xia, J. Using MetaboAnalyst 5.0 for LC–HRMS Spectra Processing, Multi-Omics Integration and Covariate Adjustment of Global Metabolomics Data. *Nat. Protoc. 2022 178* **2022**, *17* (8), 1735–1761. https://doi.org/10.1038/s41596-022-00710-w.

(11) Georgieva, G.; Badalova, M.; Feyzula, F.; Nedeva, T. Antifungal Activity of *Pseudomonas chlororaphis* 1S4. *Acta Microbiologica Bulgarica*. **2024**, *40* (03), 365–372. https://doi.org/10.59393/amb24400311

(12) Moore, D.; Robson, G. D.; Trinci, A. P. J. 21st Century Guidebook to Fungi, *2nd ed.* Section 4.9 Growth Kinetics. Cambridge University Press. **2020**. https://doi.org/10.1017/9781108776387
